# Brain X chromosome inactivation is not random and can protect from paternally inherited neurodevelopmental disease

**DOI:** 10.1101/458992

**Authors:** Eric R Szelenyi, Danielle Fisenne, Joseph E Knox, Julie A Harris, James A Gornet, Ramesh Palaniswamy, Yongsoo Kim, Kannan Umadevi Venkataraju, Pavel Osten

## Abstract

Non-random (skewed) X chromosome inactivation (XCI) in the female brain can ameliorate X-linked phenotypes, though clinical studies typically consider 80-90% skewing favoring the healthy allele as necessary for this effect^1–10^. Here we quantify for the first time whole-brain XCI at single-cell resolution and discover a preferential inactivation of paternal to maternal X at ∼60:40 ratio, which surprisingly impacts disease penetrance. In Fragile-X-syndrome mouse model, Fmr1-KO allele transmitted maternally in ∼60% brain cells causes phenotypes, but paternal transmission in ∼40% cells is unexpectedly tolerated. In the affected maternal Fmr1-KO(m)/+ mice, local XCI variability within distinct brain networks further determines sensory versus social manifestations, revealing a stochastic source of X-linked phenotypic diversity. Taken together, our data show that a modest ∼60% bias favoring the healthy allele is sufficient to ameliorate X-linked phenotypic penetrance, suggesting that conclusions of many clinical XCI studies using the 80-90% threshold should be re-evaluated. Furthermore, the paternal origin of the XCI bias points to a novel evolutionary mechanism acting to counter the higher rate of *de novo* mutations in male germiline^11–16^. Finally, the brain capacity to tolerate a major genetic lesion in ∼40% cells is also relevant for interpreting other neurodevelopmental genetic conditions, such as brain somatic mosaicism.

---

The X chromosome expresses more brain-specific genes than any other chromosome^17^ and X-linked gene mutations give rise to a high number of neurodevelopmental disorders, including Rett syndrome (RTT), Fragile X syndrome (FXS) and more than one hundred thirty X gene-linked intellectual disability and developmental disability disorders^17–23^.

In female eutherian mammals, X chromosome inactivation (XCI) is thought to be a random process by which either maternally inherited X (Xm) or paternally inherited X (Xp) is chosen for epigenetic silencing, ensuring X dosage compensation compared to males^24–27^. However, a non-random (skewed or biased) XCI pattern favoring one Xm or Xp chromosome can occur as a consequence of a stochastic XCI fluctuation or developmental selection against an X chromosome carrying a deleterious mutation. Such selection bias favoring the healthy X chromosome has been proposed to occur in unaffected or only mildly affected female carriers of X-linked brain disorders, with the degree of skewing needed to reduce phenotypic penetrance typically defined as ≥80:20 selection ratio favoring the healthy X^1–10^. The evidence for this model of XCI clinical significance, however, remains inconclusive, as some studies reported correlations between neurodevelopmental disease manifestations and XCI skewing measured in peripheral blood cells, for example, in RTT and FXS^28–35^, while others failed to identify consistent evidence to support this^6, 9, 36–41^, including three studies that directly examined XCI in postmortem brains of RTT patients instead of relying on the indirect measure of XCI in blood^6, 9, 36^. Therefore, the extent to which XCI skewing exists in normal female brain and, even more importantly, the degree of skewing necessary to influence phenotypes in heterozygous X-linked neurodevelopmental disorders remain unclear.

In the current study, we determined the degree to which X chromosome inactivation (XCI) varies between individual female mice under normal conditions and the level of XCI skewing that is necessary and sufficient to alter penetrance of X-linked neurodevelopmental phenotypes in a mouse model of FXS. We applied our sensitive methods of whole-brain imaging and computational data analyses^42–44^ to reveal a novel XCI pattern of ∼60:40 bias favoring silencing of the paternal versus maternal X chromosome, alongside a stochastic ∼20% variability across all brain regions in each individual mouse. Strikingly, this rather modest parent-of-origin bias and regional variability had clear effects on phenotypic penetrance in the FXS mouse model. First, the overall paternal bias in XCI protected offspring from the effects of the Fmr1-KO allele inherited from the father—a finding that may represent a novel evolutionary mechanism countering the damaging neurodevelopmental effects of a higher mutation rate in male germline line^11–16^. Second, the local XCI fluctuations influenced the type of behavioral phenotype observed in mice with Fmr1-KO allele inherited maternally—a result that suggests a novel stochastic source of the broad phenotypic variability seen in X-linked neurodevelopmental disorders in female patients and which can also help to localize different disease symptoms to candidate affected brain regions, which may be useful in developing therapeutic strategies.

## Systematic parent-of-origin effect on brain XCI

To obtain unbiased and complete survey of brain XCI we applied our serial two-photon tomography (STPT)-based imaging and computational methods^42–44^ to quantify maternal versus paternal active X chromosome distribution using *knock-in* Mecp2-GFP reporter mice^45–47^ in which X-linked methyl-CpG-binding protein is tagged with GFP (Mecp2-GFP), acting as a cellular reporter of the selection of the active X chromosome (Extended Data Fig. 1-2).

We first compared the total number of brain cells with maternal X active in Mecp2-GFP(m/+) mice that inherited the X-linked Mecp2-GFP allele maternally (n = 18) and the total number of brain cells with paternal X active in Mecp2-GFP(p/+) mice that inherited the Mecp2-GFP reporter allele paternally (n = 19). Surprisingly, this comparison revealed significantly more GFP+ cells in the brains of the maternal Mecp2-GFP(m/+) mice, demonstrating an average 58:42 bias towards higher paternal XCI and, consequently, an average 58:42 ratio of cells with maternal Xm active to paternal Xp active in WT brain (Fig. 1c, d; Supplementary Table 1). Notably, though, this average ∼60:40 paternal XCI bias comprised a considerable individual variability, including extreme examples of 84:16 Xm selection bias and 25:75 Xp selection bias (Fig. 1d). Therefore, stochastic variability in XCI in early development, in addition to the systematic paternal bias, also plays an important role in determining the overall Xm:Xp ratio in each brain (note that the Xm:Xp ratios were calculated by normalizing the Mecp2-GFP(m/+) and Mecp2-GFP(p/+) cell counts to the sum of Xm-active and Xp-active cell counts, which was equal to the 100% control homozygous Mecp2-GFP(m/p) cell count; Fig. 1d; See Methods).

**Fig. 1.**
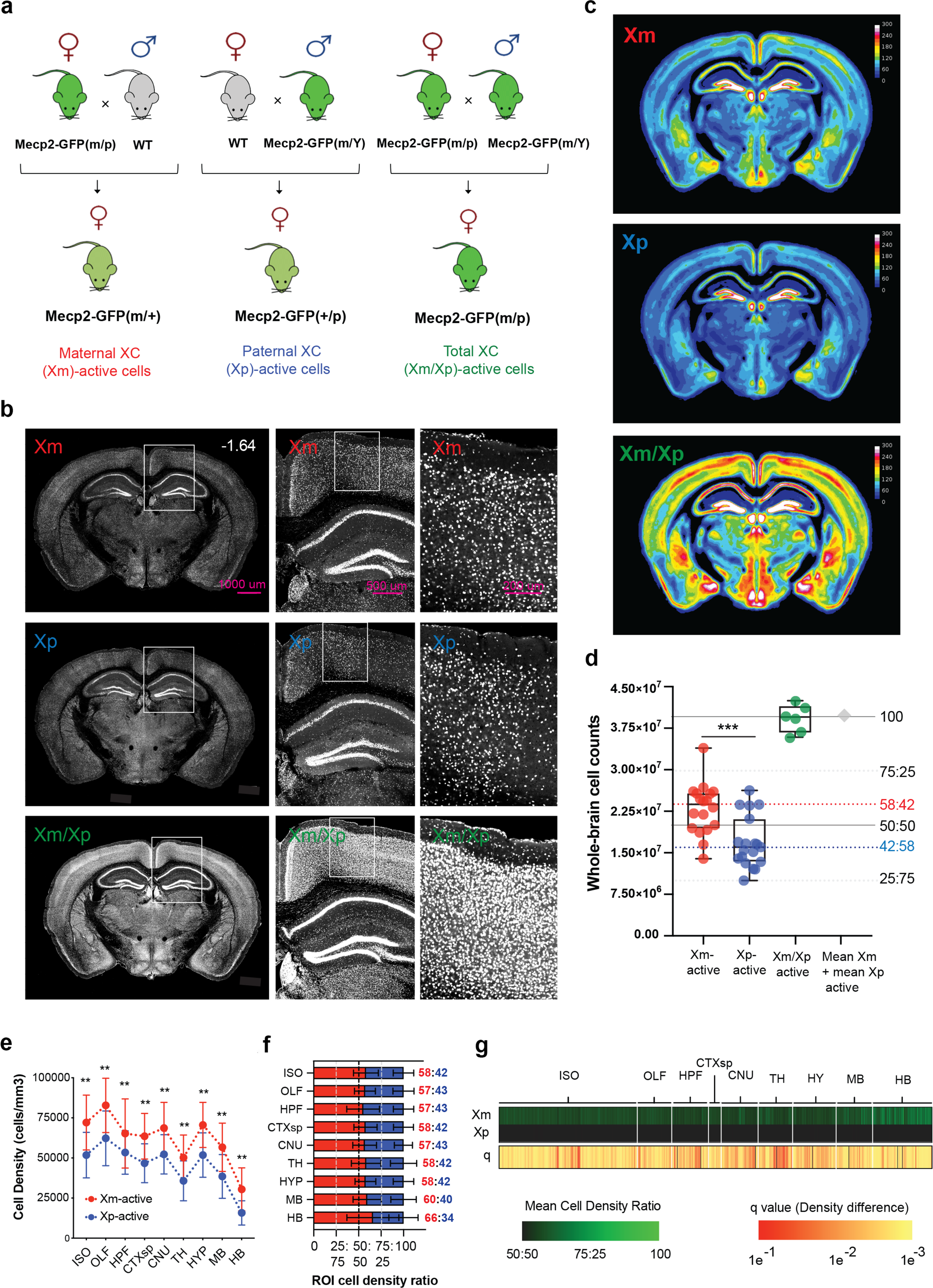
Whole-brain quantification of active maternal and paternal X chromosome distribution. **a**, Strategy for breeding maternal, paternal, and homozygous Mecp2-GFP reporter mice. Left: heterozygous Mecp2-GFP(m/+) females with maternal Mecp2-GFP allele are derived from breeding homozygous female Mecp2-GFP(m/p) and WT male mice. Middle: heterozygous Mecp2-GFP(p/+) females with paternal Mecp2-GFP allele are derived from breeding WT female and hemizygous Mecp2-GFP(m/Y) male mice. Right: homozygous Mecp2-GFP(m/p) mice are derived from breeding Mecp2-GFP(m/p) female and Mecp2-GFP(m/Y) male mice. **b**, Representative STPT images from brains with maternal Xm-Mecp2-GFP (top), paternal Xp-Mecp2-GFP (middle) and homozygous Xm-Mecp2-GFP/Xp-Mecp2-GFP (bottom) cells. **c**, Mecp2-GFP+ cell density in the three genotypes represented as voxelized heat maps of a 16-color gradient scale from black (0 cells/voxel), yellow (150 cells/voxel) to white (300 cells/100 μm sphere voxel). **d**, Whole-brain Mecp2-GFP+ cell counts (mean ± SD): Xm-GFP+ cells = 2.34 × 10^7^ ± 4.96 × 10^6^ (n=18); Xp-GFP+ cells = 1.69 × 10^7^ ± 4.63 × 10^6^ (n=19); and homozygous Xm-GFP+/Xp-GFP+ cells = 3.92 × 10^7^ ± 2.52 × 10^6^ (n=6); p = 0.00023, Welch’s t test. The sum of the heterozygous Xm-GFP+ and Xp-GFP+ cell counts is shown as a black diamond. Solid horizonal lines indicate 100% and 50% of homozygous Xm-GFP/Xp-GFP cell count. Dashed gray lines indicate 75:25 and 25:75 Xm-active:Xp-active cell ratios based on the homozygous Xm-GFP/Xp-GFP cell count. Dashed red and blue lines show the derived mean 58:42 Xm-active:Xp-active and 42:58 Xp-active:Xm-active ratios. **e**, Xm-active (red) and Xp-active (blue) cell densities (cells/mm^3^) demonstrate a comparable Xm selection bias across all major ontological brain divisions (mean ± SD; Xm-active, Xp-active): ISO (isocortex): 72118 ± 17083, 51810 ± 14188; OLF (olfactory areas): 82791 ± 16950, 62260 ± 17057; HPF (hippocampal formation): 65273 ± 21583, 53398 ± 13519; CTXsp (cortical subplate): 63541 ± 14215, 46708 ± 12138; CNU (cerebral nuclei): 68663 ± 16025, 52269 ± 12254; TH (thalamus): 50224 ± 14088, 35745 ± 12424; HY (hypothalamus): 70594 ± 14086, 51834 ± 13835; MB: 56670 ± 15046, 38513 ±1364; HB (hindbrain): 30458 ± 13392, 15766 ± 7539; **p < 0.01, 2-way mixed effects ANOVA with Holm-Sidak corrected post-hoc comparisons. **f**, Data from **e**, converted into cell density ratios reveal ∼60:40 Xm-active bias across all regions. Ratios for each ROI are listed to the right and color-coded by Xm-active (red) and Xp-active (blue) samples. **g**, Xm selection bias is seen across all brain regions. Left two columns: Heat map visualization of normalized mean Xm-active and Xp-active ROI cell density on a color gradient of black (50%) to green (100%). Right column: statistical significance across all ROIs (FDR-corrected student’s t tests), with each q value indicated by a color gradient from red (0.1) to yellow (0.001).

Next, we asked whether the paternal XCI bias seen at the whole-brain level exists similarly across all brain areas or whether there may be regional differences in XCI patterns in the brain. This analysis revealed that the average ∼60:40 bias for maternal X selection is seen across all major brain divisions, including the isocortex (58:42), cortical subplate (58:42), olfactory areas (57:43), hippocampal formation (57:43), cerebral nuclei (57:43), thalamus (58:42), hypothalamus (58:42), midbrain (60:40) and hindbrain (66:34) (Fig. 1e, f; Supplementary Table 1, 2) as well as across all local subregions in these areas (Fig. 1g; Supplementary Table 1, 2). These data thus argue against a previous model of cortical versus subcortical differences in parent of origin XCI^48^, suggesting instead that the maternal X selection bias is seen not only in the cortex but also across the entire brain (note that the anatomical segmentation of the imaged brains was done as previously described by us using the 2011 ARA mouse brain atlas;^42, 43, 49^ see Methods).

The detailed anatomical brain segmentation also allowed us to measure regional XCI variability in each individual brain (Fig. 2; Supplementary Table 1), with the aim to determine whether such variability is related to (or independent of) the overall whole-brain Xm:Xp ratio, such as whether brains with highly skewed Xm:Xp ratios comprise different (or the same) regional variability compared to brains with the overall Xm:Xp ratio close equal. To investigate this question, we visualized regional variability in Xm and Xp selection across all anatomical regions and for each brain in brain-wide 2D heatmaps (Fig. 2a) and collapsed box-and-whisker plots (Fig. 2b). This revealed a similar variability from the mean for all imaged brains independent of their overall Xm:Xp ratio with the mean coefficient of variations (CV) for both Mecp2-GFP(m/+) and Mecp2-GFP(p/+) groups being ∼20% compared to CV of ∼10% in homozygous Mecp2-GFP(m/p) mice (Fig. 2c). Taken together with the previous analyses, these data show that Xm versus Xp selection in each brain can be described as a result of a modest overall bias favoring the selection of the maternal X (and silencing of the paternal X) across all brain regions, combined with a moderate ∼20% local stochastic variability independent of the overall selection.

**Fig. 2.**
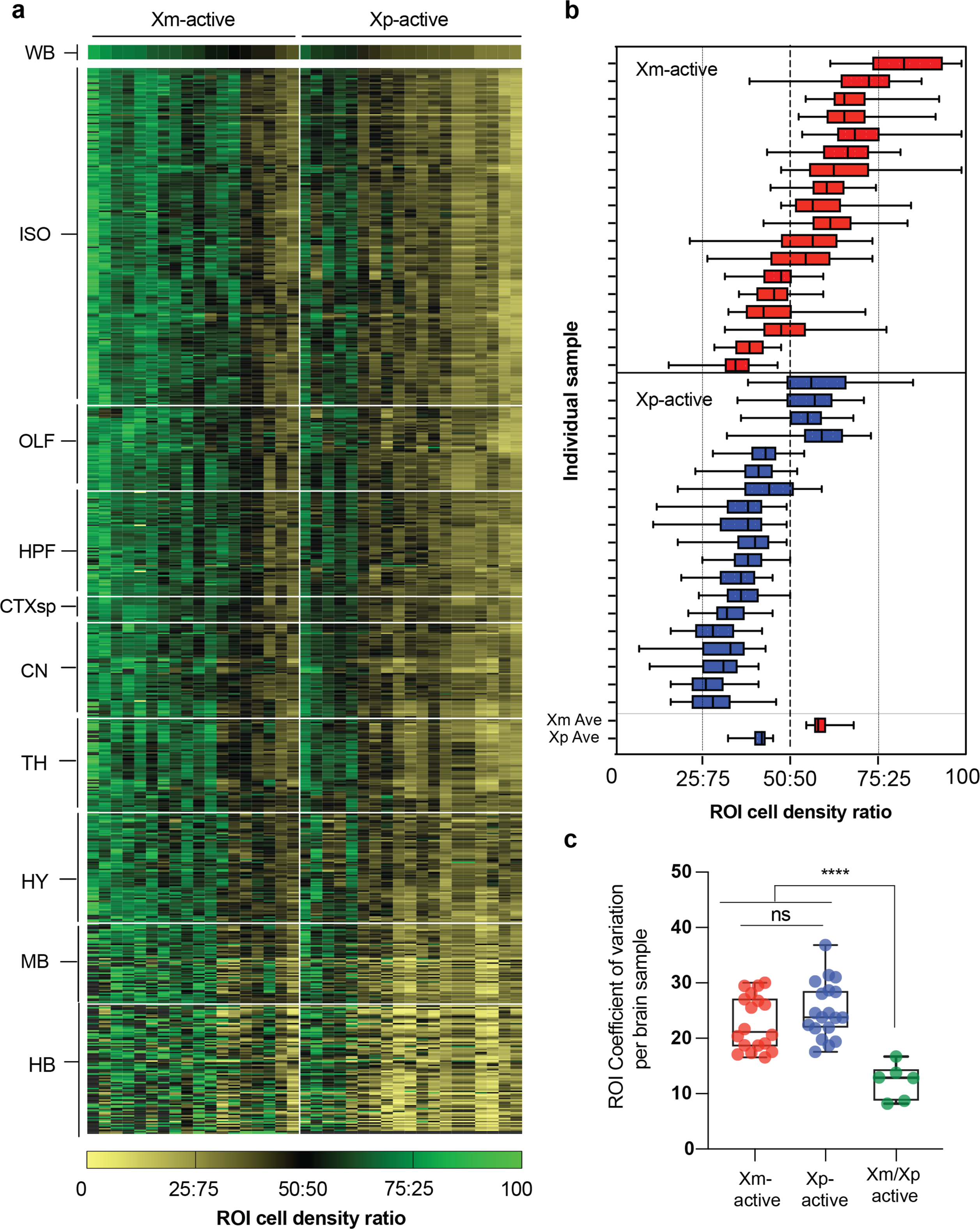
Quantification of Xm and Xp selection and its variability across all brain regions. **a**, 2D heatmap of brain-wide ROI cell density ratios in which each column represents one animal from maternal Mecp2-GFP(m/+) (left) and paternal Mecp2-GFP(p/+) (right) reporter brains. The Xm and Xp selection is expressed asmeasured:estimated cell density ratios across all 738 brain ROIs and displayed on a color gradient of beige to black (0 to 50:50), and black to green (50:50 to 100%). The samples are ordered from high (left) to low (right) XCI selection for each genetic group on the x axis whereby the ROIs are ordered by major ontological division along the y-axis. **b**, Stochastic variability of Xm versus Xp selection across all brain regions in each brain analyzed. The average data for Xm-GFP and Xp-GFP brains is plotted at the bottom. Box- and-whisker plots display median, interquartile range, and 95th percentiles of the data. **c**, Quantification of brain-wide ROI stochastic variability for Xm-GFP versus Xp-GFP versus homozygous Xm-GFP/Xp-GFP alleles by coefficient of variation analysis (CV): mean ± SD = 22.8 ± 4.9 vs 25.08 ± 5.1 vs 12.16 ± 3.2. ***p<0.005 from 1-way ANOVA with Dunnet’s post-hoc corrected multiple comparisons.; 25% percentile = 18.41 vs 21.79 vs 8.56, Median = 21.14 vs 23.78 vs 12.83, 75% percentile = 27.31 vs 28.68 vs 14.48.

## Paternal XCI bias gates FXS disease penetrance

Having quantified the XCI patterns in WT mice, we next asked whether the identified biases may affect disease penetrance in a female heterozygous mouse model of the fragile X syndrome (FXS), an X-linked disorder caused by the loss of expression of the Fragile X mental retardation 1 (Fmr1) protein^18, 50^. Based on the WT data, we envisioned the following scenarios: First, the average ∼60:40 bias favoring maternal X selection should lead to more brain cells carrying the mutant allele inherited from the mother than from the father, resulting in more pronounced phenotypes associated with the maternal transmission. Second, stochastic variability in regional Xm:Xp selection may further modify the phenotypic outcomes based on which brain areas comprise the least favorable mutant-to-healthy X ratios. On the other hand, a positive selection for cells with the healthy X chromosome may change the XCI distribution from what is seen in WT brain, resulting in lesser than expected skewing and related phenotypes.

To test these scenarios, we crossed the Fmr1 knock-out (KO) mouse model of FXS^51^ with the Mecp2-GFP X reporter line, generating heterozygous Fmr1-KO/+ female mice with the KO allele inherited either maternally in Fmr1-KO(m)/Mecp2-GFP(p) mice or paternally in Fmr1-KO(p)/Mecp2-GFP(m) mice (Fig. 3a). We note that while this FXS mouse model has been studied extensively as a complete knock-out in male hemizygous Fmr-1 KO/Y mice, only three studies reported modest phenotypes (synaptic and social) in female heterozygous Fmr1 KO/+ mice with the KO allele always transmitted maternally^52–54^. Hence, our study is the first to directly compare the phenotypic effect of maternal versus paternal Fmr1 KO allele transmission in heterozygous female mice, while also quantifying the whole-brain distribution of the Fmr1-KO allele-expressing cells in each animal.

**Fig. 3.**
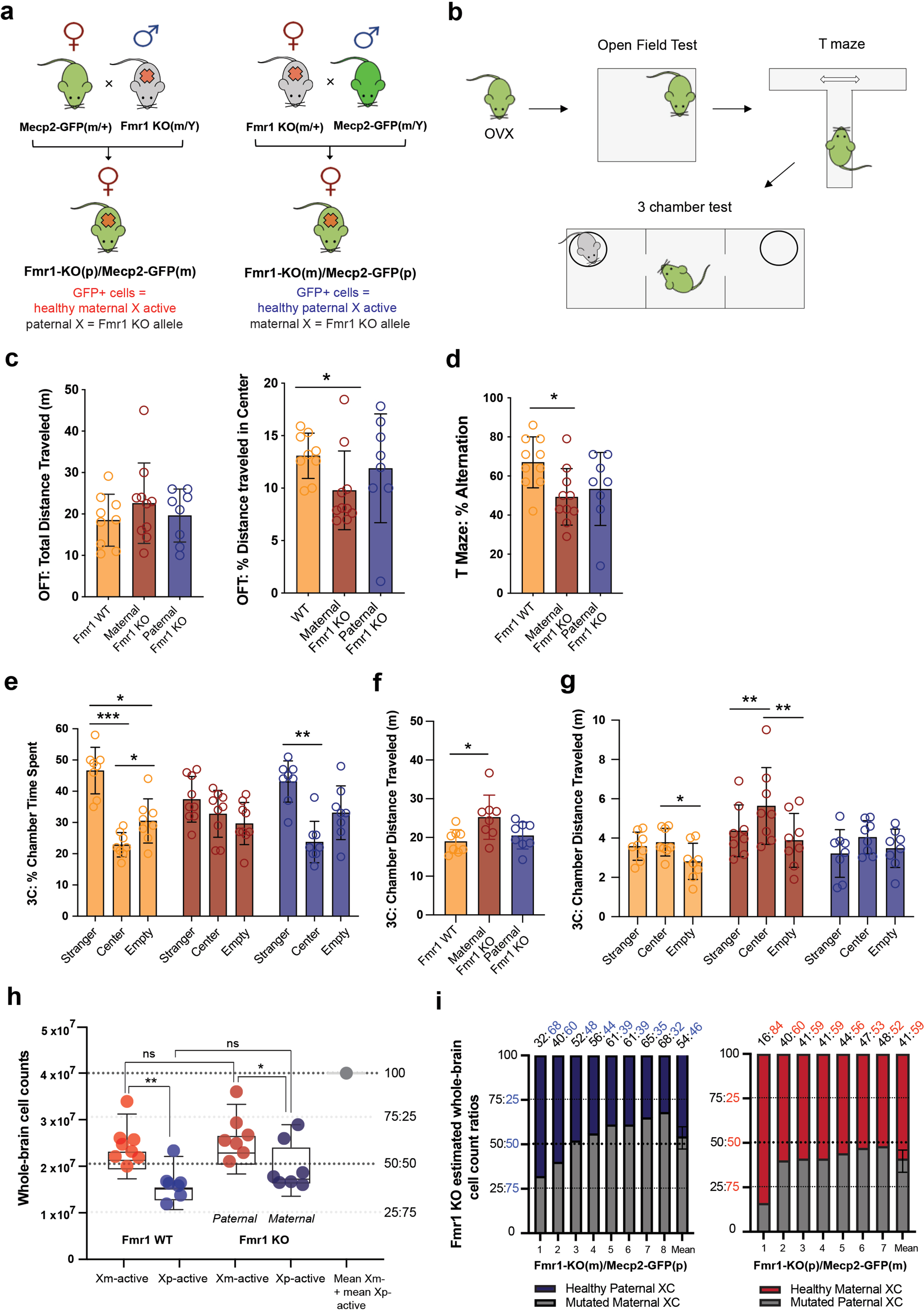
Maternal transmission of the Fmr1-KO allele at an average 46:54 Fmr1-WT:KO cell ratio is sufficient for phenotypic penetrance. **a**, The Fmr1 KO allele was transmitted either maternally in Fmr1-KO(m)/Mecp2-GFP(p) mice (in figures for short Fmr1-KO(m)/+; left) or paternally in Fmr1-KO(p)/Mecp2-GFP(m) mice (in figures Fmr1-KO(p)/+; right) by breeding with Mecp2-GFP male or female mice, respectively. **b**, Behavioral testing: ovariectomized (OVX) heterozygous maternal Fmr1-KO(m)/+, paternal Fmr1-KO(p)/+ and WT littermate mice were sequentially tested in the open field test (OFT), T maze, and 3-chamber social test (n=8 for each group). Data for each individual test are derived from sequential data of the same animals. **c**, OFT results show reduced exploration only in maternal Fmr1-KO(m/+) mice: Left: Total distance traveled was not different among the three groups (mean ± SD; meters): WT: 19.2 ± 6.3; Fmr1-KO(m/+): 23.2 ± 10.6; Fmr1-KO(+/p): 19.6 ± 6.3. Right: Percent distance traveled in center arena was reduced in Fmr1-KO(m)/+ mice (mean ± SD): WT: 13 ± 2; KO(m/+): 9 ± 3; KO(+/p): 12 ± 5. ANOVA, K-W = 6.47; p = 0.039; Welch’s ANOVA and Dunnet T3 post-hoc test: WT vs Fmr1-KO(m/+): p = 0.035. **d**, T maze results show reduced working memory, seen in reduced percent spatial alterations, only in maternal Fmr1-KO(m/+) mice (mean ± SD): WT: 69 ± 14; Fmr1-KO(m/+): 47 ± 15; Fmr1-KO(+/p): 53 ± 19. Welch’s ANOVA, Dunnet T3 post-hoc comparisons: WT vs Fmr1-KO(m/+): p = 0.023. **e**-**g**, 3-chamber test identifies deficits only in maternal Fmr1-KO(m/+) mice. **e**, Fmr1-KO(m)/+ show a lack of sociability in the 3-chamber test. Time spent for the 3 genotypes in social vs center vs empty chamber were (mean ± SD): WT = 47 ± 7 vs 23 ± 4 vs 31 ± 7; Fmr1-KO(m/+) = 39 ± 6 vs 32 ± 8 vs 29 ± 7; Fmr1-KO(+/p) = 43 ± 7 vs 24 ± 7 vs 33 ± 9; 2-way mixed effects ANOVA, Holm-Sidak post-hoc comparisons; *p<0.05; **p<0.01; ***p<0.001. **f**, Fmr1-KO(m)/+ mice were hyperactive compared to WT mice as seen in increased total distance traveled (mean ± SD): 24.25 ± 6.10, 18.96 ± 3.07 and 20.49 ± 3.48, respectively. Welch’s ANOVA and Dunnet T3 post-hoc comparisons, WT vs Fmr1-KO(m/+): p = 0.037. **g**, The hyperactivity of the Fmr1-KO(m)/+ mice was restricted to the starting central arena of the 3-chamber apparatus. Distance traveled for the 3 genotypes in social vs center vs empty chamber were (mean ± SD): WT = vs 3.79 ± 0.71 vs 3.08 ± 1.17; Fmr1-KO(m)/+ = 4.37 ± 1.32 vs 5.63 ± 1.96 vs 3.80 ± 1.36; Fmr1-KO(p)/+ = 3.22 ± 1.21 vs 4.05 ± 0.84 vs 3.71 ± 0.98; *p<.05; **p<.01, 2-way mixed effects ANOVA, Holm-Sidak post-hoc comparisons. **h**, Quantification of whole-brain Xm vs Xp selection in WT mice (left) and heterozygous Fmr1 KO mice (right) (mean ± SD): WT: Xm-active cells = 2.6 × 10^7^ ± 4.7 × 10^6^ and Xp-active cells = 1.7 × 10^7^ ± 3.6 × 10^6^; maternal KO allele in Fmr1-KO(m/+): healthy Xp-active cells in = 2.0 × 10^7^ ± 5.2 and paternal KO allele in Fmr1-KO(p/+): healthy Xm-active cells = 2.6 × 10^7^ ± 4.8. *p<0.05, **p<0.01, ANOVA with Holm-Sidak post-hoc comparisons. Estimated 75:25 and 25:75 whole-brain cell count ratios are shown in dashed lines. **i**, Stacked bar graphs of each maternal Fmr1-KO(m/+) (left) and paternal Fmr1-KO(+/p) (right) whole-brain X selection from (**h)**: healthy Xp-active cells in Fmr1-KO(m)/+ brains are highlighted in dark blue color (left) and healthy Xm-active cells in Fmr1-KO(p/+) are highlighted in red color (right). Fmr1-KO:WT cellular ratios of each sample is listed above.

We first assayed the impact of the maternal versus paternal Fmr1-KO allele transmission across three behavioral tests that were used previously to identify disease-related phenotypes in FXS mice: the open field (OFT) test to assess sensorimotor functions and anxiety-related behavior^55–61^; the T-maze spontaneous alternation test to assess working memory^60, 62, 63^; and finally the 3-chamber test to assay sociability and social preference^61, 64, 65^ (Fig. 3b). Strikingly, these experiments revealed that while the Fmr1-KO(m)/Mecp2-GFP(p) heterozygous female mice with maternal KO allele transmission showed deficits in all three tests, whereas the paternal Fmr1-KO(p)/Mecp2-GFP(m) heterozygous mice were not different from control Mecp2-GFP sibling mice in any measurement (Fig. 3c-g).

The behavioral deficits of the maternal Fmr1-KO(m)/Mecp2-GFP(p) mice included: 1) reduced travel distance across the center arena in the OFT suggesting reduced exploratory behavior due to sensorimotor deficits and/or increased anxiety (Fig. 3c), 2) reduced frequency of spontaneous alterations in the T-maze suggesting impaired working memory (Fig. 3d), and 3) a complete lack of social preference in the 3-chamber social interaction test, reflected by no preference for time spent in the chamber with a stranger mouse presented under a wire cup versus the control chamber comprising only an empty cup (Fig. 3e-g). Interestingly, the maternal Fmr1-KO(m)/Mecp2-GFP(p) mice were also hyperactive in the 3-chamber apparatus, as reflected by an increased total distance traveled compared to control Mecp2-GFP sibling mice, and this hyperactivity was restricted to the starting middle chamber (Fig. 3f, g). This suggests a more complex social phenotype of a combined hyperactivity and avoidance of both social and non-social stimuli in Fmr1-KO(m)/Mecp2-GFP(p) mice.

Taken together, our behavioral results demonstrate that female mice with the Fmr1 KO allele transmitted maternally, but not paternally, display disease-related phenotypes. This, in turn, suggests that the bias of higher maternal X selection (and higher paternal X silencing) seen in WT mice persists in the heterozygous Fmr1-KO/+ mice and results in more brain cells carrying the Fmr1 KO allele inherited maternally than paternally. To test this prediction directly, we imaged the brains of all mice used in the above behavioral tests by STPT as done for WT brains in Fig. 1 and determined the distribution of the WT Fmr1 allele-expressing cells marked by Mecp2-GFP expression from the same X chromosome. These measurements revealed the following. First, the whole-brain Xm:Xp ratio in the maternal Fmr1-KO(m)/Mecp2-GFP(p) mice was 54:46, representing an average 54% of Fmr1-KO allele expressing brain cells compared to 46% Fmr1 WT allele expressing cells (Fig. 3h, i; Supplementary Table 3, 4). Second, the regional Xm:Xp ratio differences were more pronounced for cortical versus subcortical areas (Extended Data Fig. 3), suggesting a modest compensation favoring the selection of the healthy paternal Xp chromosome subcortically compared to WT brains. And third, the whole-brain Xm:Xp ratio in the paternal Fmr1-KO(p)/Mecp2-GFP(m) mice was 41:59, reflecting an average 41% of Fmr1-KO allele expressing brain cells compared to 59% Fmr1 WT allele expressing cells (Fig. 3h-i). These data thus demonstrate that an average ∼55:45 mutant to healthy brain cell density ratio is sufficient to cause behavioral phenotypes in the maternal Fmr1-KO(m)/Mecp2-GFP(p) mice, while in contrast ∼40:60 mutant to healthy brain cell density ratio is below a threshold needed to produce phenotypic manifestations in the paternal Fmr1-KO(p)/Mecp2-GFP(m) mice.

## Local XCI variability contributes to phenotypic diversity

Brain-wide cellular distribution measurements of Fmr1 WT versus KO alleles allowed us to further test whether regional stochastic XCI variability may further influence the observed phenotypes in the affected maternal Fmr1-KO(m)/Mecp2-GFP(p) mice. For example, it may be expected that mice with high Fmr1-KO allele cell density in brain areas known to regulate social behavior will show more pronounced phenotypes in the 3-chamber sociability test compared to those with a stochastically lower Fmr1 KO allele cell density in the same brain areas. To test this hypothesis, we correlated the Fmr1 WT:KO cell ratios across all brain regions to behavioral scores from all three behaviors for each mouse tested. This correlation analysis identified two distinct sets of anatomical regions in the maternal Fmr1-KO(m)/Mecp2-GFP(p) brains, in which the Fmr1 WT:KO cell ratios were indeed correlated to behavioral performance in either the OFT or the 3-chamber test (Fig. 4a; Extended Data Fig. 4).

**Figure 4.**
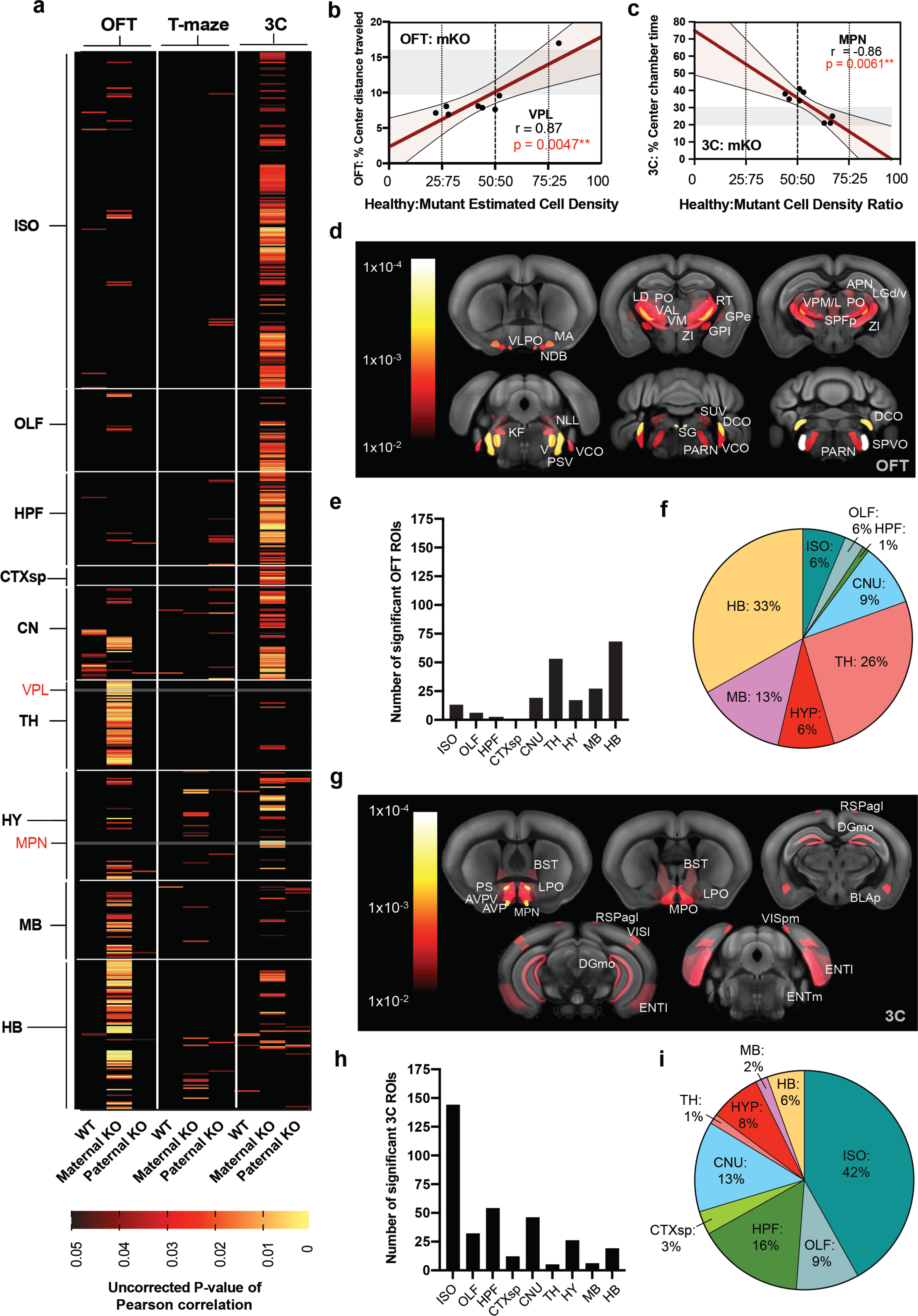
Identification of two sets of brain regions with local Fmr1-WT:KO cell ratios linked to phenotypes in the OFT and 3-chamber test. **a**, Identified brain regions in Fmr1-KO(m)/Mecp2-GFP(p) (“Maternal KO”) mice that correlate Fmr1 WT:KO allele ratios to behavioral performance in the OFT and 3-chamber tests. Data are displayed as 2D heat maps of statistically significant correlations across brain ROIs grouped by major ontological structures. Uncorrected P values of Pearson correlations are represented on a color gradient scale from 0.05 (black) to 0.005 (red) to 0.0005 (yellow). Results from Fmr1 WT (“WT”), maternal Fmr1-KO(m)/Mecp2-GFP(p) (“Maternal KO”) and paternal Fmr1-KO(p)/Mecp2-GFP(m) (“Paternal KO”) mice are grouped by each behavioral test: left: OFT, center: T maze, right: 3-chamber (3C). **b-c**, Representative scatterplot displays of correlated ROI density from (**a**) shown for (**b**) VPL of thalamus in OFT and (**c**) MPN of hypothalamus in 3 chamber tests. **d**, **g**, Selected ROIs from maternal Fmr1-KO(m)/Mecp2-GFP(p)\ mice with significant correlation (cut-off at p=0.01) to performance in (**d**) OFT and (**g**) 3 chamber tests are heat mapped and overlaid on a reference mouse brain template. **e**, **h**, Number of significantly correlated ROIs from maternal Fmr1-KO(m)/Mecp2-GFP(p) mice listed by major brain structure for (**e**) OFT and (**h**) 3 chamber tests. **f**, **i**, 100% pie charts of significantly correlated ROIs from maternal Fmr1-KO(m)/Mecp2-GFP(p) mice grouped by major brain structure and represented as a percent of total correlated ROIs for (**f**) OFT and (**g**) 3-chamber test. ROI acronyms: VLPO – ventrolateral preoptic area; NDB – nucleus of the diagonal band; MA – magnocellular nucleus; PO – posterior complex of the thalamus; VPM/L – ventral posteromedial/lateral nucleus of the thalamus; PC – paracentral nucleus; RT – reticular nucleus of the thalamus; ZI – zona incerta; GPi – globus pallidus, internal; GPe – globus pallidus, external; SPFp – subparafacsicular nucleus, parvicellular part; LGv – ventral part of the lateral geniculate nucleus; LGd – dorsal part of the lateral geniculate nucleus; KF – Koelliker-Fuse subnucleus; V – motor nucleus of the trigeminal; PSV – principal sensory nucleus of the trigeminal; VCO – ventral cochlear nucleus; SG – supragenual nucleus; DCO – dorsal cochlear nucleus; VCO – ventral cochlear nucleus; SPVO – spinal nucleus of the trigeminal, oral part, E, F) PS – parastrial nucleus; BST – bed nuclei of the stria terminalis; AVP – anteroventral preoptic nucleus; AVPV – anteroventral periventricular nucleus; MPO – medial preoptic nucleus; LPO – lateral preoptic area; RSPAgl – retrosplenial cortex, agranular layer; DGmo – dentate gyrus, molecular layer; BLAp – basolateral amygdala, posterior; VISpm – posteromedial visual area; VISl lateral visual area; ENTl – entorhinal area, lateral; ENTm – entorhinal area, medial.

The first set of brain regions comprised areas in which the Fmr1 WT:KO cell ratio was positively correlated to OFT behavioral performance, specifically the distance traveled in the center of the OFT arena. These regions included primarily sensory structures of the thalamus, midbrain and hindbrain (Fig. 4a, d-f; Extended Data Fig 4a; Supplementary Table 5), such as the sensory ventral posterolateral nucleus of the thalamus (VPL) (Fig 4b). These data thus indicate that a higher density of cells expressing the Fmr1 WT allele across the identified areas known to regulate sensorimotor functions, among others, translates to increased exploration of the open field arena and hence a reduced OFT phenotype. The second set of brain regions comprised areas in which the Fmr1 WT:KO cell ratio was inversely correlated to the time spent in the center of the 3-chamber apparatus. These regions, in contrast to the first set, contained primarily cortical, hippocampal and hypothalamic brain areas (Fig. 4a, c, g-i; Extended Data Fig. 4a; Supplementary Table 5), including the hypothalamic medial preoptic nucleus (MPN) that is well known for regulating social behavior (Fig 4c; Extended Data Fig 4c). These data, in turn, indicate that a higher density of cells expressing the Fmr1 KO allele across areas known to regulate social behaviors, among others, translates to more time spent in the starting chamber of the 3-chamber apparatus and hence a more pronounced social deficit phenotype. The same correlation analysis failed to identify a distinct set of brain regions with Fmr1 WT:KO ratios related to behavioral performance in the T-maze test in which the Fmr1-KO(m)/Mecp2-GFP(p) mice showed only a modest level of impairment (Fig. 3a; Supplementary Table 5). In addition, whole-brain Fmr1 WT:KO ratios, in contrast to the regional ratios described above, showed only a trend towards a positive correlation in the OFT and negative correlation in the 3-chamber task (Extended Data Fig. 5). This suggests that while the whole-brain WT:KO cell ratios set the overall risk for disease penetrance, the regional Fmr1 WT:KO cell ratios determine the specific behavioral phenotypic outcome in each animal. Finally, as expected, we also did not observe any significant correlations for brain regions in the paternal Fmr1-KO(p)/Mecp2-GFP(m) mice which did not show any behavioral phenotypes (Fig. 4a; Supplementary Table 5).

**Fig. 5.**
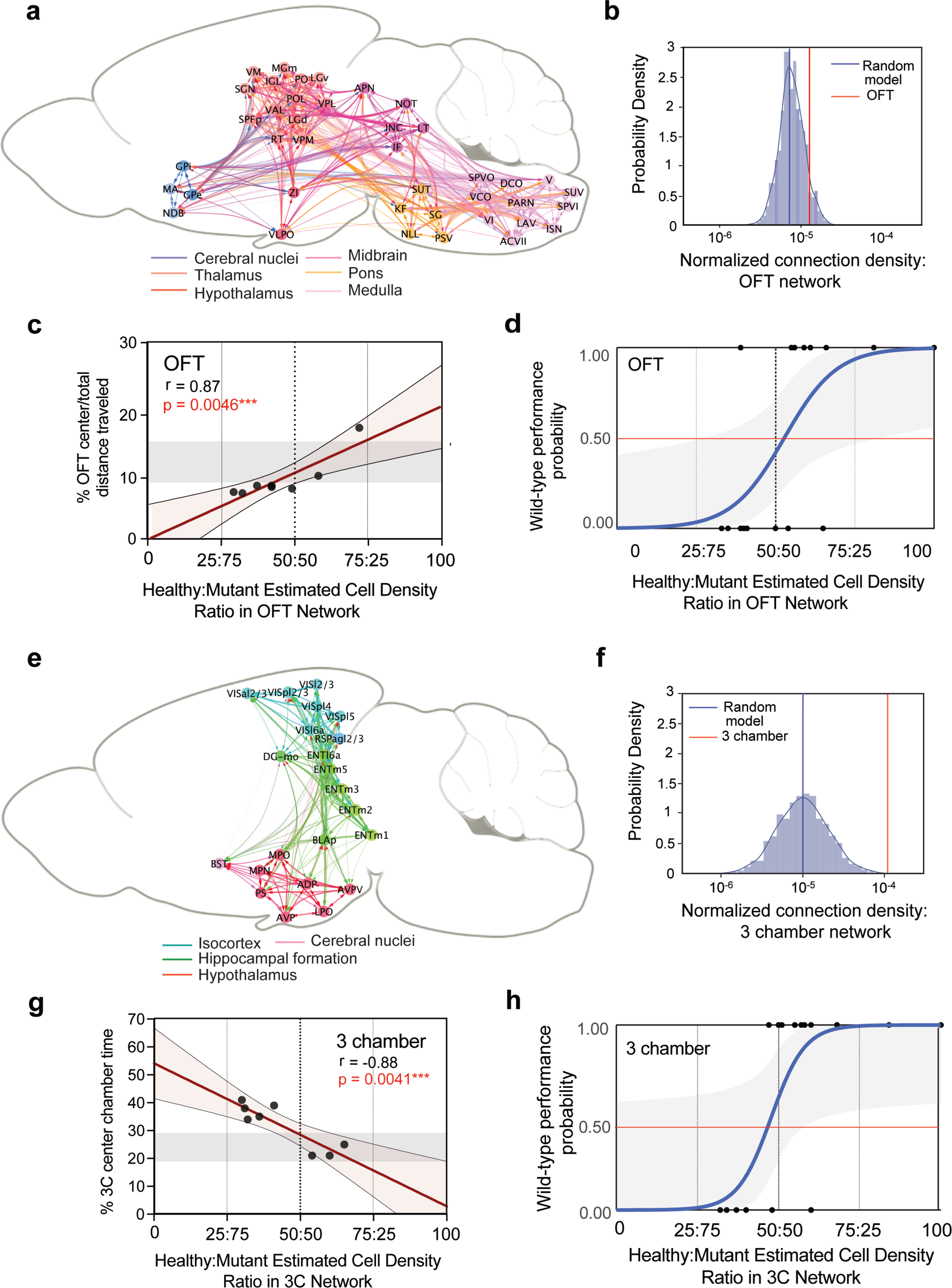
FXS phenotypic penetrance in the OFT and 3-chamber tests is determined by XCI-defined healthy:mutant cellular ratios within structural brain networks. **a**, **e**, Visualization of structural connectivity weights within brain networks of ROIs with WT:KO allele ratios correlated to behavioral performance in the (**a**) OFT and (**e**) 3-chamber tests (See Methods). **b**, **f**, Normalized median connection densities of the (**b**) OFT (1.22e^-6^) and (**f**) 3-chamber (9.95e^-4^) ROI networks (red lines) overlaid on the probability density of 1000 in-silico generated random ROI networks of the same inter-regional distance and ROI amount (blue line = median; OFT: 7.22 x10^-6^; 3 chamber: 9.95 × 10^-6^). One sample t test: OFT: p = 6.36 x 20^-247^; 3-chamber: p = 0.0). Data is compared and presented on a log scale. **c**, **g**, Linear regression models of behavioral scores and ROI network cell density ratios in (**c**) OFT and (**g**) 3-chamber assays. Regression and statistical test values are listed for each panel. Shadowed rectangle in each group represents control Fmr1 WT range of behavioral scores for comparison. **d**, **h**, Logistic regression modeling of WT behavioral performance predicted by healthy cell density percent in ROI networks of behavioral penetrance from all mice (n=22) within (**d**) OFT (K; *x*^2^(1) = 14.88; log-likelihood ratio = -14.42; equal odd ratio = 54%; p = 0.00011) and (**h**) 3 chamber (L; *x*^2^(1) = 13.88; log-likelihood ratio = -12.89; equal odd ratio = 49%; p = 0.00019) behaviors. Grayed area represents ± 95% confidence intervals.

## Anatomical brain networks underlying distinct behavioral phenotypes

The identification of the two sets of brain regions with Fmr1 KO allele density linked to behavioral phenotypes suggests that these represent two distinct anatomically connected brain networks regulating sensorimotor and anxiety-related versus social behaviors. To test this hypothesis further, we next applied the recently established structural connectivity matrix analysis derived from a whole-brain connectivity model of the mouse brain^66^ (See methods). This analysis indeed revealed much higher connection densities for brain regions within each brain network than for matching randomly sampled brain structures: the OFT network density (Fig. 5a-b) and 3-chamber network density (Fig. 5e-f) represented the 93^rd^ and 100^th^ percentile of each sample network’s distribution, respectively (Supplementary Table 6; see Methods). This analysis thus supports the model where the brain areas with Fmr1-KO cell density linked to different behavioral deficits represent two distinct functional brain networks.

The identified correlations of local Fmr1-KO cell densities to behavioral phenotypes also suggest that the distribution of the Fmr1-KO allele across the two brain networks determines and can in fact predict the disease penetrance in each animal. To test this hypothesis, we next calculated the Fmr1 WT:KO allele ratios selectively across the two brain networks and regressed these values against the behavioral performance in the OFT and 3-chamber test (Supplementary Table 7). As shown in Fig. 5, the Fmr1 WT:KO allele ratios were indeed highly significant predictors of individual behavioral performance in only maternal Fmr1-KO(m)/Mecp2-GFP(p) heterozygous mice in both the OFT (Fig. 5c) and 3-chamber (Fig. 5g) assays, and not for control or paternal Frm1-KO(p)/Mecp2-GFP(m) mice (Extended Data Fig. 6. These data thus further support the model where the cellular distribution of the Fmr1-KO allele across the two brain networks represent the cellular and genetic substrate the respective OFT and 3-chamber test behavioral phenotypes.

Finally, in the last set of analyses, we asked what is the local Fmr1 KO cell density that can differentiate normal from disease-related behavior. Likelihood-ratio tests performed on binary logistic regression models revealed that the Fmr1-KO allele distributions across the brain networks indeed significantly predicts normal from disease-related performance in both the OFT (Fig. 5d) and 3-chamber test (Fig. 5h). The equal-odds ratio of normal versus disease-related behavioral outcome was calculated to be 55.20 ± 5.95% healthy cell density percent in the OFT brain network and 49.18 ± 5.19% healthy cell density percent in the 3-chamber brain network. This phenotypic penetrance threshold of ∼50% regional Fmr1-KO cell density agrees with the lack of behavioral manifestations when the Fmr1-KO allele is transmitted paternally in ∼40% brain cells (Fig 3).

## Discussion

We applied our automated whole-brain imaging platform^42^ to comprehensively quantify the respective distributions of active maternal and paternal X chromosomes across the entire mouse brain at single cell resolution. This approach allowed us to discover subtle yet both statistically and functionally significant patterns that were missed by previous studies relying on semiquantitative observations typically from only a few selected brain areas^48, 67–69^.

Clinical studies of the effects of XCI skewing on neurodevelopmental disorders typically consider >80% (or even >90%) skewing in favor of the healthy X chromosome as significant with respect to modifying X-linked phenotypes, with ratios below the 80% threshold defined as balanced (i.e., non-skewed)^1–10^. Therefore, these studies suggest that brain development can tolerate only 10 to 20% of brain cells carrying a deleterious mutation and a higher percentage of affected brain cells should be expected to lead to clinical manifestations. In contrast, our study revealed a lack of behavioral phenotypes in paternal heterozygous Fmr1-KO(p)/+ mice carrying the Fmr1-KO allele in ∼40% brain cells, demonstrating an unexpectedly robust brain capacity to compensate for the presence of a harmful mutation in nearly half of brain cells. Therefore, we propose that a threshold of ∼60% skewing in favor of the healthy X chromosome may be sufficient for modifying neurodevelopmental phenotypic penetrance and should be considered clinically relevant. This suggests that negative results from many clinical studies that considered >80% XCI skewing as a threshold for interpretating clinical phenotypes may need to be reevaluated with the lower threshold >60% as a skewed XCI definition^5, 37–40, 70–76^.

In addition to challenging the dogma that only large degree of XCI skewing can affect X-linked neurodevelopmental phenotypes, our results also show that stochastic XCI variability, estimated from our measurements to be ∼20% across all brain regions independent of the overall brain XCI bias, can play an important role in determining the specific phenotypic manifestations in each individual female. Our data revealed that in the maternal heterozygous Fmr1-KO(m)/+ mice, the Fmr1-KO allele variations across two distinct brain networks can predict the likelihood of disease-related phenotypes in two behavioral tests – the open field and 3-chamber test measuring sensorimotor performance and sociability, respectively – in each animal. Thus, while the overall ∼60% brain cell transmission of the maternal Fmr1-KO allele predisposes for disease related phenotypes, the ∼20% stochastic brain-wide variation on top of the overall pattern is likely to determine the specific features of the phenotypic representation (Fig 6). This finding provides an important etiological insight into the potential source of phenotypic variability in X-linked neurodevelopmental syndromes disease in female patients.

**Fig. 6.**
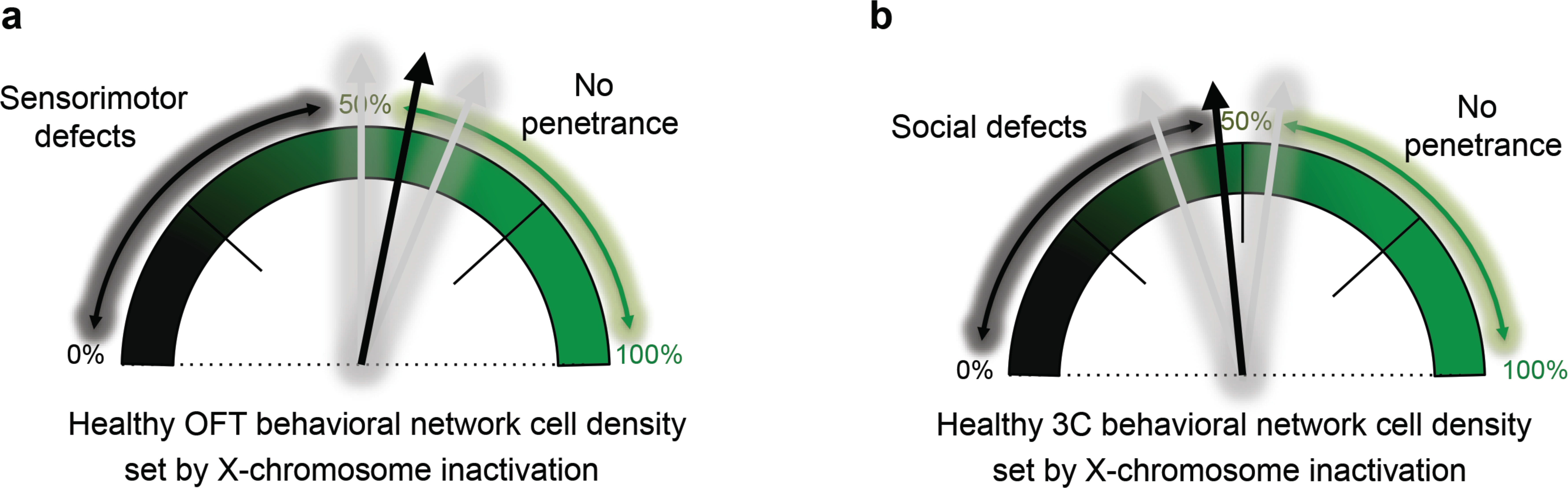
Cartoon depiction model of (**a**) OFT and (**b**) 3 chamber logistic modeling results (Fig 5), which portrays how female FXS phenotypes are determined by the distributed healthy cell density occupying the ROI behavioral networks identified in this study.

The finding that the brain can tolerate the loss of as important a gene as Fmr1 in nearly half brain cells may also be relevant for the interpretation of the significance of somatic mutations in the brain, which occur during cell divisions after fertilization and lead to a mosaic expression of a genetic lesion in a subset of brain cells and cell types depending on the cell lineages affected^77–80^. While some genetic lesions can lead to neurodevelopmental phenotypes even when expressed in a relatively local cell population in the brain, such mutations linked to focal epilepsy or gross morphological development^81–,87^, our data suggest that mutations related to more cognitive dysfunctions in neurodevelopment, such as those linked to autism or intellectual disability, may require a similarly high threshold of >50% brain cells (perhaps within a specific cell type lineage) to be affected to lead to phenotypic manifestations.

Finally, our results also suggest a novel evolutionary role of the observed 60:40 XCI paternal bias. Maternally and paternally imprinted autosomal genes are expressed in highly distinct regional patterns proposed to mediate competing interests of each parent in terms of the evolutionary fitness of their genes, as proposed in the so-called kinship or genetic conflict theory^88, 89^. Our study revealed a systematic ∼60:40 paternal bias in XCI across all brain divisions and regions, suggesting that XCI is not part of such a maternal versus paternal genetic competition. On the other hand, the paternal XCI bias may play a different evolutionary role. It is well established that the mutation rate in the paternal germline is significantly higher compared to that in the maternal germ cells, which should result in a higher rate of X-linked mutations linked to neurodevelopmental deficits from paternal transmission^11–16^. Our findings thus suggest that the selective bias to silence ∼60% of the paternal X chromosome and the surprising developmental capacity to tolerate the mutant allele in nearly half brain cells may in fact represent a novel evolutionary mechanism acting to counter the higher frequency of *de novo* mutations inherited from the father compared to the mother in daughters.

In the current study we used the Fmr1-KO allele as an established mouse model of X-linked genetic lesion, but we note that human FXS is caused by expansion (>200 repeats) of the 5’ UTR CGG sequence of an FMR1 premutation allele inherited exclusively maternally^18, 90–92^. Therefore, the observed ∼60:40 paternal XCI bias is not directly pertinent to FXS inheritance, but instead may be evolutionary important in protecting offspring from a higher rate of paternally inherited *de novo* mutations in one of the more than 130 X genes linked to non-syndromic intellectual disability and other neurodevelopmental phenotypes^17–23^.

## Supplemental Tables

Table 1: Brain-wide descriptive statistics of cell counts, volumes, and cell densities

Table 2: Brain-wide statistical results of Xm-active versus Xp-active cell density comparisons

Table 3. Brain-wide descriptive statistics of cell counts, volumes, and cell densities

Table 4. Statistical results of brain-wide cell density comparisons in Fmr1 WT or heterozygous KO mice

Table 5: Statistical results of brain-wide ROI cell density and behavioral score correlational screens

Table 6: Normalized ROI connection density matrices

Table 7: ROI networks of penetrance raw data

## Methods

### Animal breeding and husbandry

Adult mice (8-10 weeks old) were used for whole-brain imaging experiments. Animals were housed under a 12-hour light/dark cycle (0600 ON, 1800 OFF), had access to food and water ad libitum, and were housed with littermates. All experimental procedures were performed in accordance with CSHL Animal Care and Use Committee Guidelines. The Mecp2-GFP mouse line was obtained from the Jackson laboratory (stock # 014610). Mecp2 is a gene located at chromosomal position X A7.3 and is subject to XCI. Developed in the laboratory of Adrian Bird, this mouse line contains an in-frame knock-in cassette at the 3’ UTR of the Mecp2 locus ^46, 47, 93, 94^. Driven and regulated by the endogenous Mecp2 promoter/enhancers, Mecp2-GFP expression leads to normal Mecp2 levels and subcellular localization of Mecp2 protein that is fused at the C-terminus with EGFP. Expression of the fusion allele does not alter neuronal physiology ^94^ and mice are successfully bred to homozygosity without behavioral or reproductive complications (data not shown). In addition, strong expression of Mecp2-GFP favors neurons of many types ^46^ thereby circumventing biased effects of XCI determinations based on expression profile. Mecp2-GFP(m/+) or Mecp2-GFP(p/+) mice were obtained in separate heterozygotes by crossing homozygous females or hemizygous males with wild-type C57Bl6/J (JAX stock # 000664) mice. A subset of these wild type reporter mice was derived from Fmr1 KO or WT crosses that generated Mecp2-GFP(m/+)/Fmr1 KO(+/+) (n=7) and Mecp2-GFP(+/p)/Fmr1 KO(+/+) (n=8) mice that are congenic to the C57Bl6/J crosses. Homozygous reporter mice were obtained by crossing homozygous Mecp2-GFP(m/p) females with hemizygous Mecp2-GFP(m/Y) males. Fmr1 KO mice were obtained from the Jackson laboratory (#003025). These mice were originally developed in the Oostra laboratory and contain a gene-disrupting neomycin resistance cassette in exon 5 of the FMR1 locus ^51^. Mecp2-GFP(m/+)/Fmr1 KO(+/p) female mice were generated by breeding Mecp2-GFP(m/p) females with hemizygous Fmr1 KO(m/Y) males. For imaging only, Mecp2-GFP(m/+)/Fmr1 KO(+/+) female mice were generated by separately breeding homozygous Mecp2-GFP(m/p) females with hemizygous Fmr1 KO(+/Y) males. Conversely, Mecp2-GFP(+/p)/Fmr1 KO(m/+) or Mecp2-GFP(+/p)/Fmr1 KO(+/+) wild type littermates were generated by breeding Fmr1 KO(m/+) females with hemizygous Mecp2-GFP(m/Y) males. Using this genetic strategy, double heterozygous mice used for behavior and imaging experiments contained the Mecp2-GFP and Fmr1 KO alleles on opposing X chromosomes. All transgenic mice were maintained on a C57Bl6/J background.

### Brain sample preparation

Animals were euthanized via transcardial perfusion under ketamine/dexmedetomidine anesthesia. Dissected brains were post-fixed overnight in 4% paraformaldehyde at 4 C, incubated for 48 h in 0.1 M glycine/0.1 M PB for auto fluorescent quenching, and then stored in 0.05 M PB at 4 C until confocal or serial two-photon tomography imaging (STPT; see below). Prior to STPT imaging, brains were embedded 4% oxidized agarose in 0.05 M PB using custom molds and holders to maintain consistent embedding position. Embedded brains were crosslinked in 0.2% sodium borohydrate solution for 3h at room temperature or overnight at 4 C prior to STPT processing (below).

### Immunohistochemistry and confocal imaging

Neuronal expression of the MeCP2-GFP allele was studied though immunostaining and confocal imaging. 50 um vibratome-processed, free-floating coronal sections of homozygous MeCP2-GFP mice brains (n=2) were processed. Sections were washed 3 times in PBS followed by blocking for 1 h at room temperature in PBS-T (PBS, 0.2% Triton-X 100) containing 5% donkey serum. Sections were then incubated overnight at 4 C in blocking solution containing rabbit anti-NeuN (Millipore, ABN78) primary antibody at 1:1000. After washing, NeuN-stained sections were incubated with anti-rabbit AlexaFluor-568-conjugated secondary antibody (Thermo-Scientific, A10042) diluted 1:500 for 1 h at room temperature. After washing excess secondary antibody, sections were mounted, DAPI-counterstained (Prolong Gold Antifade Mountant, Thermo Fisher), and coverslipped for imaging. Confocal images were acquired with a Zeiss LSM780 confocal microscope using a 561 laser and corresponding dichroic and filter sets. Single plane images were captured with a 40x oil immersion objective. Total colocalized populations for each marker of every FOV (212.55 um X x 212.55 um Y) were manually quantified using Fiji image processing package.

### Serial two-photon tomography whole-brain image acquisition

The Tissuecyte1000 instrument was used for all imaging experiments (TissueVision) ^44^. This system combines a high-speed multi-photon microscope with a fully integrated vibratome for automated z-sectioning and image acquisition throughout the entire whole-mount sample. Embedded sample brains were imaged with a 20x objective at 50 μM below the sample surface. 270 total serial sections were acquired at 50 um z-resolution (13.5 mm total z-length), with each section being comprised of a 12 (x-axis, 700 um) × 16 (y-axis, 700 um) field of view (FOV) mosaic. Images were acquired with laser scan settings of 1 um/pixel at an integration time of 1 us. A laser wavelength of 910 nm with ∼322 mW power at the end of the objective was used for optimal excitation/emission of MeCP2-GFP fluorescence. Constant laser settings and PMT detector settings were used for all samples.

### Automated MeCP2-GFP+ cellular detection and counting

Raw image tiles for each brain were illumination corrected, stitched in 2D with Matlab and aligned in 3D using Fiji software ^44^. For reliable automated MeCP2-GFP detection from full brain datasets, we implemented convolutional networks (CNs) ^95^. CN training for detection of MeCP2-GFP+ cells in the STPT datasets was accomplished as in previous studies ^43^ with CN training performed on human marked-up ground truth data (biological expert identified MeCP2-GFP+ nuclei) of MeCP2-GFP brains. CN performance was determined based on F-score calculations (F-score = the harmonic mean of the precision and recall, where precision is the ratio of correctly predicted cells divided by all predicted cells and recall is ratio of correctly predicted cells divided by ground truth positive cells; ∼1800 MeCP2-GFP+ cells were marked/expert/brain). Composite F-scores for MeCP2-GFP CN was obtained by determining F-scores in 8 FOVs (400 (X) um by 400 (Y) um) representing different cellular density and imaging content in 3 separate heterozygous MeCP2-GFP+ brains (24 FOVs total). Stable precision and recall was seen for all regions analyzed, delivering a composite F-score of 0.84 (Extended data Fig. 2). In the CN output images, signal smaller than 10 μm^2^ was removed as noise. In order to normalize the performance of CN for each brain, the brightness of MeCP2-GFP+ signal for each sample was normalized by the mean and standard deviation of tissue autofluorescence signal from a coronal section corresponding to bregma position of +0.20 mm. We did not analyze MeCP2-GFP+ cells in the cerebellum due to faulty brain-to-brain warping of this region (data not shown).

### 3D brain registration and anatomical segmentation

Registration of individual brains to a standardized reference space was computationally achieved as published previously ^43^. In short, affine transform was calculated using 4 resolution levels and B-spline with 3. Advanced Mattes mutual information ^96^ was the metric used to measure similarity between moving and fixed images. Image similarity function is estimated and minimized for a set of randomly chosen samples with each 23 images in a multi-resolution and iterative fashion^44^. Entire warping of whole-brain images is done using Elastix ^97^. Anatomical segmentation of Allen Brain Atlas (ABA) labels onto sample brains was made possible also as previously published ^43^. Version 2.2 ABA labels (836 total) were transformed onto individually registered samples. Quality control of ROI segmentation found and excluded 95 ROIs total from analysis due to erroneous counting most likely caused by small ROI size and/or warping location (Full ROI list found in Supplementary Table 1). In addition, cell counts from layer 6 a and b were combined into one layer, layer a.

### 2D-3D cell count correction and density measurements

Detected 2D cell count values obtained at 50 um Z resolution were transformed by a stereological 3D conversion factor obtained by the following way (Extended data Fig. 2). First, counting boxes of 200 um x 200 um x 50 um (xyz) were acquired at 2.5 um Z resolution via optical imaging within 6 brain regions comprising major anatomical divisions of a female heterozygous MeCP2-GFP mouse brain. 20 optical images were acquired at a depth range that spanned 50 um around the normal 50 um focal depth (i.e. 25-75 um below the tissue surface). Second, Mecp2-GFP CN was run on the middle optical section corresponding to the 50 um depth. Third, manual markup of Mecp2-GFP+ nuclei was performed in each counting box using the stereological counting rules of Williams and Rakic ^98^. Lastly, a conversion factor for each region was calculated by dividing manual 3D counts by 2D CN count of the middle section. This factor was averaged over the 6 regions reaching a final conversion factor of 2.6. (Extended data Fig. 2b). ROI cellular density was obtained by 1) transforming ABA labels onto individual brains, 2) converting ROI assigned pixel space to mm^3^, 3) dividing 2.5 um Z-corrected absolute cell counts by mm^3^ values by to arrive at cells/mm3.

### Behavioral testing

6-8 month old ovariectomized female mice were behaviorally phenotyped in a sequential series of tests. All mice were ovariectomized at least 2 weeks prior to testing in order to remove estrous cycle influences from behavior. Each behavioral test was separated by 2-7 days to avoid acute post-testing and handling effects. Mecp2-GFP(+/p)/Fmr1 KO(+/+) mice served as behavioral controls for all behaviors studied. The following tests were sequentially performed on each mouse:

#### Open field test (OFT)

To measure activity and anxiety in an open field, unhabituated mice were placed in a 40 x 40 x 40 cm2 open plexiglass box containing a layer of fresh bedding. The open field arena was located in a non-sound-proof, enclosed environment under dim lighting. All mice were housed in the same facility room behavioral testing was performed. An overhead camera visually captured all tests and ANY-maze (Stoelting) automated behavior tracking software was used for real-time activity/location recording and analysis. A 20 x 20 cm center square designated within the tracking settings defined the center and perimeter boundaries of the arena. The software measured total and center distance traveled. For center-specific activity, center distance was normalized to total distance traveled and presented as percent total distance traveled. Adequate cleaning of the maze with bleach, water and drying was performed between each mouse. Fresh bedding was added to the arena for each subject.

#### T-maze

We studied mouse spatial memory by measuring spontaneous spatial alternations in the T-maze ^99, 100^. Spontaneous alternation is an innate exploratory behavior possessed by rodents which is hippocampus-dependent and serves as an index of spatial and working memory ^99^. Our protocol was based off of the continuous version with minor modification^100^. The dimensions of the T-maze used was 35 cm stem length, 28 cm arm length, 10 cm arm height, and 5 cm lane width (Stoelting). For testing, the T-maze was located in a non-sound-proof, enclosed environment under dim lighting. All mice were housed in the same facility room behavioral testing was performed in. To begin the test, each mouse was carefully placed at the stem start position of the maze and was freely allowed to enter either arm. To prevent the mouse from entering the other arm after its initial choice, a metal block was placed at the entrance of the empty arm once the subject committed exploration to an arm. The subjects were allowed to freely explore the chosen arm and stem until it explored back to start of the stem. Once the beginning position was reached, the mouse was held in-between the start position and a metal block placed proximally to the start position for 5 seconds. The metal block was then removed and the mouse was allowed again to enter an arm of its choice. Manual scoring of each arm choice and time to experimental completion was made after 14 trials. No more than 3 minutes/trial was allowed for each subject and encouragement was given to each subject at 3 minutes (in the form of hand movement behind the mouse) to return to start position. Mice that did not complete more than 9 trials were excluded from analysis. Adequate cleaning of the maze with bleach, water and drying was performed between each mouse. The number of trial-to-trial arm entry alternations (e.g. left-to-right or right-to-left) was calculated and expressed as a percent of total trials.

#### 3-chamber test

Sociability was measured using the 3-chamber test based on the protocol developed in the Crawley laboratory^102^. The 3-chamber apparatus used consisted of a plexiglass box (60 x 40 x 22(h) cm) partitioned into 3 chambers (20 cm/each) (Stoelting). Doors (4 x 8 cm) connecting chambers allowed the mice to freely explore all areas of the box. The apparatus was located in a non-sound-proof, enclosed environment under dim lighting. All mice were housed in the same facility room that behavioral testing was performed in. An overhead camera visually captured all test sessions and ANY-maze (Stoelting) automated behavior tracking software was used for real-time activity/location recording and analysis. Chamber designations in tracking software were user-defined and used for chamber-specific activity measurements. Two metal-barred cylindrical cages (7 cm (diameter) × 15 cm (height); 3 mm bar diameter and 7 mm spacing) were used for stranger mouse containment in one chamber and for an empty enclosure in the opposite-sided chamber. The cage bars are spaced such that close sniffing is the only interaction type possible. Ovariectomized adult female Fmr1 WT mice were used as stranger mice and were habituated to an enclosure cage for 10 minutes at least 1 day prior to any experiments. Each stranger mouse (n=8) was used 4 times only and were rotated every 4 experiments for use. Test mice were habituated to an empty 3 chamber apparatus for 10 minutes prior to actual experiments. For testing, mice were allowed to freely explore all chambers for 10 minutes. For each experiment the enclosed stranger mouse was placed in the left chamber and the empty enclosure on the right. Chamber time spent and distance traveled was quantified for each chamber. Percent time spent or distance traveled was calculated as total value/individual chamber value.

### Quantification of structural connectivity within correlated brain networks of ROIs

We determined if OFT and 3-chamber significantly correlated ROI groups (herein referred to as “networks”) represented structural connectivity networks by comparing the median ROI network connection weight to a distribution of randomly sampled networks of the same size for both tasks. ROIs having significant correlation of p<0.01 were included in the analysis. We used the normalized connection density, a measure of connection strength normalized by both source and target region sizes, from the regional structural connectivity matrix ^66^. We restricted the population of structures for each network to ROIs from which we could draw to an intermediate level of the ontology represented by 292 ’summary structures’ ^101^. The intersection of these summary structures with the sets of ROIs for each task resulted in sets of 39 and 13 ROIs for the open field and 3-chamber tasks, respectively (**Table S6**). These ROIs included: OFT: ACVII, APN, DCO, GPe, GPi, IF, IGL, ISN, LAV, LGd, LGv, LT, MA, MG, NDB, NLL, NOT, PARN, PB, PO, POL, PSV, RT, SG, SGN, SPFp, SPVI, SPVO, SUT, SUV, V, VAL, VCO, VI, VLPO, VM, VPL, VPM, ZI; 3 chamber: ADP, AVP, AVPV, BST, ENTl, ENTm, LC, LPO, MPN, MPO, PS, VISl, VISpl. Additionally, since mesoscale connectivity is distance dependent ^101^ and the model in ^66^ is spatially dependent, we restricted the selection of random ROI networks to have similar inter-regional distance dependence as that of their respective cell density correlated ROI networks.

The procedure for this selection is as follows:

Given a set of N cell density correlated summary structure level ROIs:

1. Randomly draw a set of N regions from the set of summary structures.
2. Compute the pairwise inter-regional distances for the set of sampled regions.
3. Compute the Kolmogorov-Smirnov statistic to measure the difference in distributions of distances for the sampled and cell-count correlated networks

a. If the KS statistic shows a significant difference in distributions (having a p-value < 0.01), reject the sample and return to (1).
b. Else, return the sample.

The above procedure is repeated 1000 times, after which the median normalized connection density of the experimental ROI networks is compared to the distribution of the sample medians. Since these connectivity measures are log-normally distributed ^66, 101^ the t statistic is computed in log-transformed space to test the significance of the difference. Visualization of ROI-ROI connectivity for each behavioral network of ROIs was created with Cytoscape network visualization and analytic program (Version 3.7.1). Significantly correlated ROIs of the deepest ontological distance from root structures were chosen for visualization, except for the ROIs not annotated at the summary structure level and hence not found in the structural connectivity matrix. Those ROIs included: BSTmg, BSTpr, BSTif, BSTpr, PVHap, TTv3, isl, islm, MPNc, MPNl, PAA3, and COApl3. Log-transformed normalized connection densities were used for edge sizes and scaled for presentation. Edges with sizes < 0 were excluded from the visualization.

### Statistics

Whole-brain absolute cell counts were compared amongst Xm-active and Xp-active reporter brains using a Welch’s t test. A 2-way mixed effects ANOVA ([2] XC-active/KO or WT allele parent-of-origin x [9] major ROI) was used to compare XC-active (Fig. 1) or WT:KO allele (Fig. 2) parent-of-origin across cell densities from major ROIs. Holm-Sidak post-hoc tests were used to assess simple between-subjects effects. Brain-wide screens for skewed XCI were statistically performed using Benjamini-Hochberg FDR-corrected student’s t tests on XC-active cell density groups. Welch’s ANOVAs with Dunnet’s T3 post-hoc testing was used to compare group performances in the OFT, T maze, and 3 chamber total distance traveled. Chamber by group 3 chamber results were analyzed with 2-way mixed effects ANOVAs with Holm-Sidak post-hoc tests of between- and/or within-subjects simple effects. Whole-brain absolute healthy cell counts in Fmr1 WT and KO mice were analyzed via ANOVA with Holm-Sidak post-hoc comparisons. Whole-brain cell count data and behavioral scores were correlated with Pearson’s correlation. Correlational screens were used to localize the physical source of behavioral penetrance using Pearson’s correlation amongst cell density and behavioral score across 736 ROIs. In this analysis, we did not correct the p-values against Type I error risk in favor of revealing ROI networks or patterns that share behavioral dependencies. Additionally, noise correlations (Supplementary Table 5) in the control WT groups did not surpass 5% of ROIs in both OFT (20/736 ROIs = 2.7%) and 3 chamber (5/736 ROIs = 0.6%) screens, further supporting the use of uncorrected p values in this dataset. ROI networks of behavioral penetrance were defined by p<0.01 bins of significance and each bin’s distributed cell count, volume, and density across each OFT and 3 chamber networks were calculated (Supplementary Table 7) and used for linear and logistic regression modeling. Logistic regression was performed on ROI network healthy cell density percent as the continuous, independent variable and WT performance as the categorical, dependent variable. Mice (n=22) from all genotypes were categorized as WT or mutant performers for each test based on the performance range of Fmr1 WT mice. Fmr1 WT mice were coded as containing 100% healthy cell density. A likelihood ratio test was performed on each logistic model to determine statistical significance. All statistical testing was performed with Graphpad Prism software version 7.0 and R (R Core Team). Alpha level was set at 0.05 in all analyses except where otherwise noted above.

## Acknowledgements

We would like to thank CSHL Hillside animal husbandry services for their support and efforts, members of the Osten lab for inputs on the study and Dr. Kristin Baldwin for comments on the manuscript. This work was funded by grants R01 MH096946 and U01 MH105971 to P.O and funds from The Gertrude and Louis Feil Family Trust to PO.

## Author Contributions

E.R.S. and P.O. conceptualized the study. E.R.S., P.O., and Y.K. designed the imaging experiments. R.P. performed genotyping and animal husbandry and assisted with experimental design. E.R.S. and K.U.V. implemented CN algorithms for automated cell detection. E.R.S. designed and D.F. performed all behavioral experiments. E.R.S. performed tissue processing and imaging experiments and most data analyses. J.K. and J.A.H. designed and performed brain network structural connectivity weight analyses. J.A.G. performed logistic regression analyses. E.R.S. and P.O. wrote the manuscript.

## Declaration of Interests

The authors declare no competing interests.

## Extended Data Figure and Figure Legends

**Extended Data Fig. 1.**
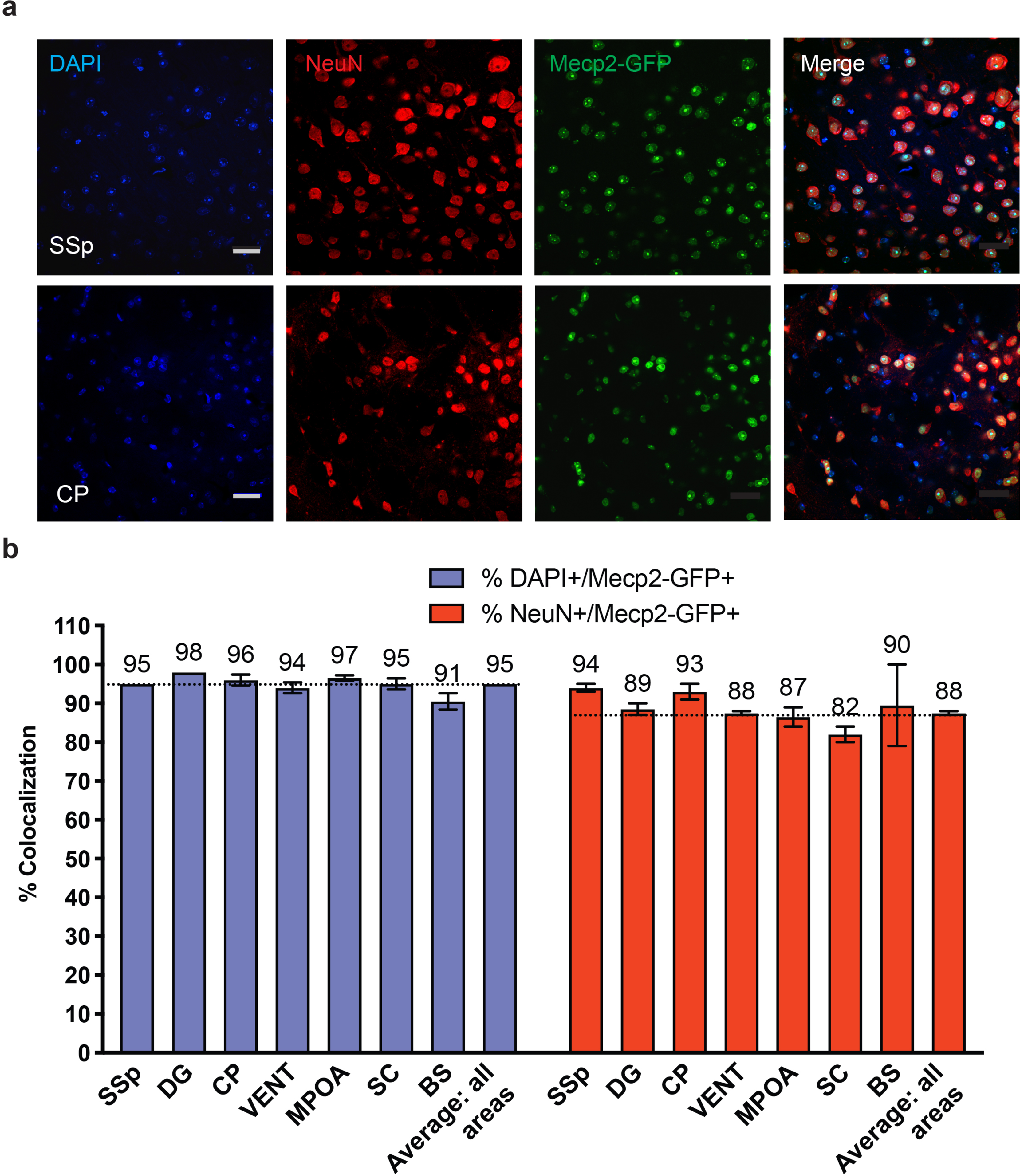
Mecp2-GFP allele labels nearly all cells in homozygous Mecp2-GFP(m/p) brain. **a**, Representative images of DAPI and NeuN counter-stained sections from somatosensory cortex (SSp) and caudate putamen (CP) areas in a homozygous Mecp2-GFP(m/p) brain (scale bar = 25 um). **b**) Quantification of DAPI and NeuN colocalization with Mecp2-GFP(m/p) expression across seven brain areas (mean ± SD): SSp = 95 ± 0 and 94 ± 1.41; DG (dentate gyrus) = 98 ± 0 and 89 ± 2.12; CP = 96 ± 1.41 and 93 ± 2.83; VENT (ventral group of the thalamus) = 94 ± 1.41 and 88 ± 0.71; MPOA (medial preoptic area) = 97 ± 0.71 and 87 ± 3.54; SC (superior colliculus) = 95 ± 1.41 and 82 ± 2.83; BS (brain stem) = 91 ± 2.12 and 90 ± 14.85 from 2 Mecp2-GFP(m/p) brains. Mean across all areas = 95 ± 0 and 88 ± 0.71.

**Extended Data Fig. 2.**
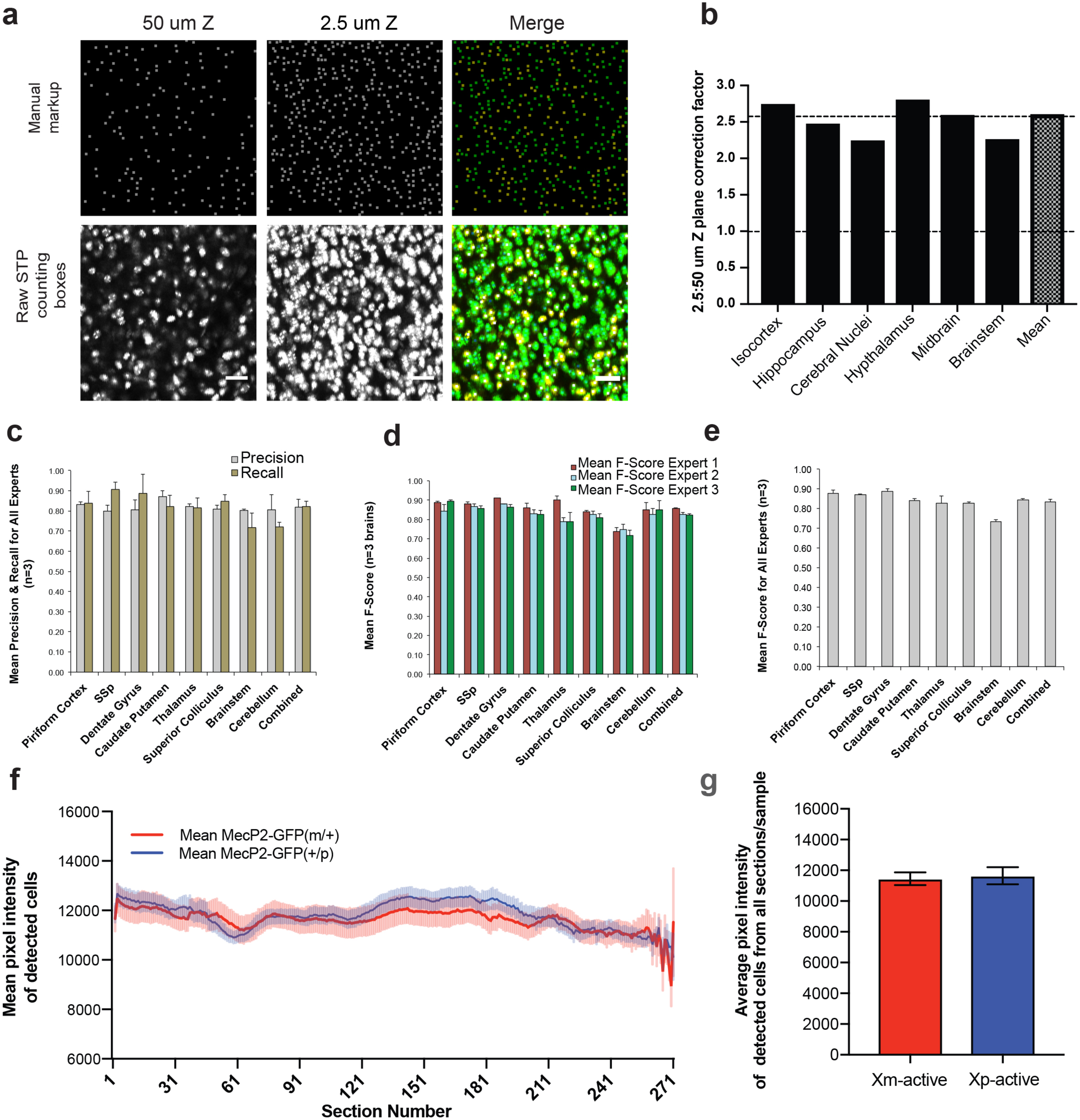
Validation and benchmarking of Mecp2-GFP+ cell detection. **a-b**, Data used to calculate a serial 2D to 3D cell count conversion factor. **a**, An example of a manual cell count markup (top row) and raw STPT image (bottom row) from a homozygous Mecp2-GFP(m/p) brain (ventromedial hypothalamus). Manual markup was made for a 3D brain volume imaged at a Z resolution of 2.5 μm. The 3D manual cell count was compared to a convolutional network (CN) based cell count from a single plane in the middle of the 3D stack. Scale bar = 25 um. **b**) 2D to 3D conversion factors were calculated by dividing the manual 3D cell counts by the single plane CN-based cell count for selected brain areas. The mean conversion factor of 2.6 was then used for all brain regions. **c**-**e**) F-score calculations of CN performance in detecting GFP+ cells based on expert ground truth data (see Methods). The calculations were performed on 8 select tiles from different brain regions covering a range of cell densities from 3 Mecp2-GFP(+/p) brain samples. **a**) Mean CN precision and recall. **b**) Mean CN F-score derived from precision and recall values of (c) for each individual expert across regions. C) Mean CN F-score shown for all regions. **f**-**g**) Fluorescence intensity comparisons of Mecp2-GFP+ cell nuclei across genotypes. **f**) Mean cellular pixel intensity from each section (270 sections total; anterior limit = 1; posterior limit = 270) of Xm-active (red) or Xp-active (blue) cells. **g**) Group comparison of mean pixel intensity across all segmented cells. All values = mean ± SEM.

**Extended Data Fig. 3.**
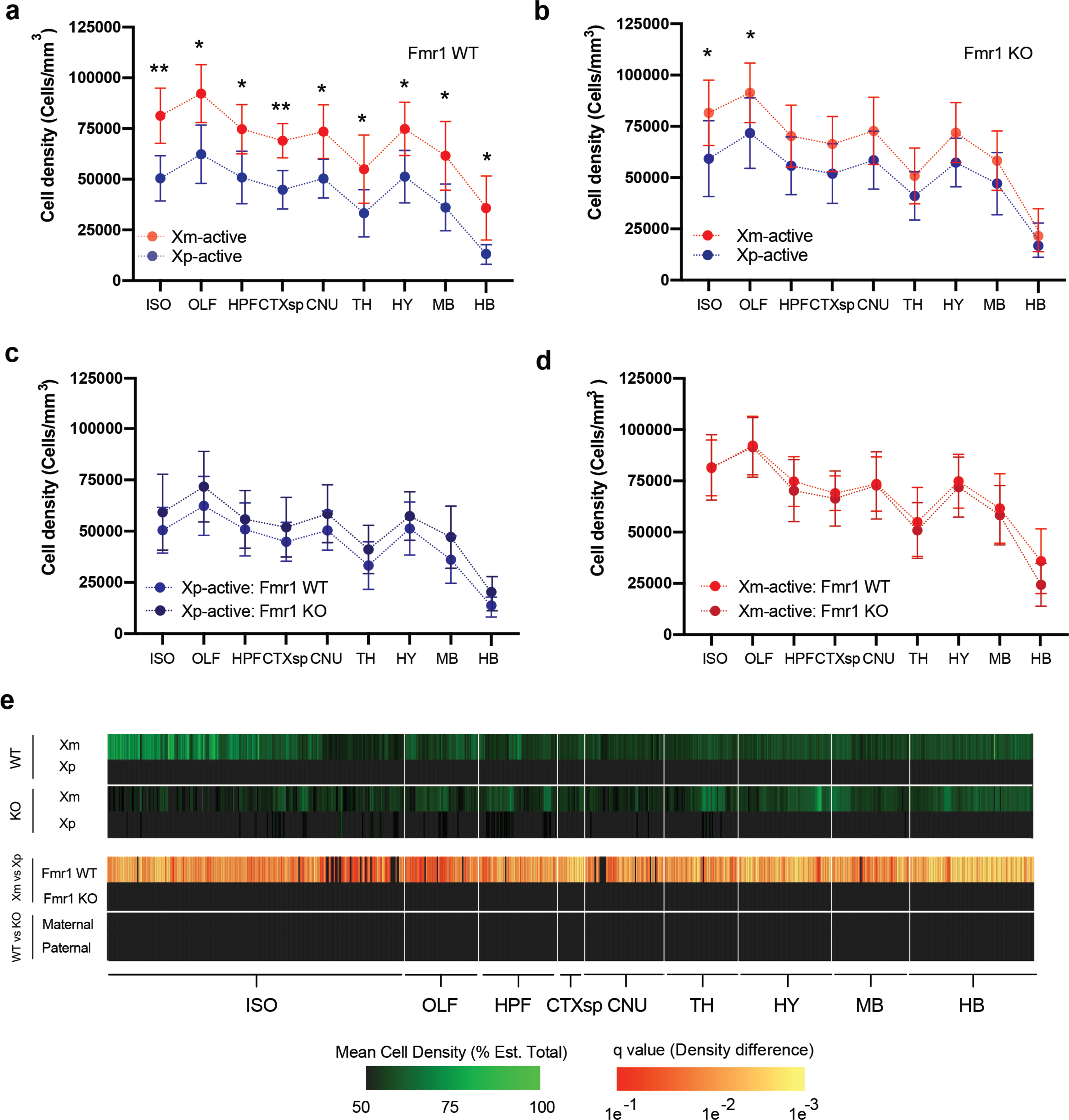
ROI-based healthy cell density analyses in Fmr1 WT and heterozygous KO mice. **a-d**, Healthy cell density (cells/mm^3^; mean ± SD) comparisons amongst (**a**) Xm-active and Xp-active Fmr1 WT, (**b**) Xm-active and Xp-active Fmr1 KO, (**c**) Xp-active Fmr1 WT and KO, and (**d**) Xm-active Fmr1 WT and KO mice across major ontological divisions of the brain. Fmr1 WT (Xm-active (n=8), Xp-active (n=7): ISO: 81354 ± 13598; 50458 ± 11153; OLF: 92208 ± 14279, 62357 ± 14412; HPF: 74715 ± 12112, 50902 ± 12888; CTXsp: 68989 ± 8458, 44848 ± 9479; CNU: 73486 ± 113216, 50317 ± 9551; TH: 54995 ± 16794, 33252 ± 11610; HY: 74840 ± 13100, 51308 ± 12890; MB: 61564 ± 16908, 36108 ±11524; HB: 35855 ± 15759 versus 12918 ± 4819) or heterozygous KO mice (Xm-active (n=7), Xp-active (n=8): ISO: 83006 ± 15294, 54469 ± 13584; OLF: 92689 ± 13965, 67429 ± 13076; HPF: 71677 ± 14524, 52120 ± 10280; CTXsp: 67988 ± 13240, 48068 ± 10198; CNU: 74039 ± 15614, 55089 ± 10992; TH: 51358 ± 12678, 39105 ± 11187; HY: 72572 ± 13680, 54570 ± 9459; MB: 60011 ± 14278, 43524 ± 12182; HB: 25308 ± 10080, 17737 ± 7177) *p<0.05, **p<0.01, 2-way mixed effects ANOVA with Holm-Sidak multiple comparison correction. **e**) Brain-wide cell density comparisons in Fmr1 WT and heterozygous KO mice. Columns 1-4: Brain-wide heat map visualization of mean healthy Xm-active and Xp-active ROI cell densities (% of estimated total) on a color gradient of black (50%) to green (100%) amongst Fmr1 WT and KO mice. Columns 5-8: Brain-wide q values of ROI cell density statistical comparisons of active XC parent-of-origin amongst Fmr1 WT and Fmr1 KO mice (top), and cell density comparisons amongst mice with WT or KO Fmr1 alleles with matched active XC parent-of-origin (bottom). Legends indicating color scaling for mean cell density (left) and q values (right) are listed at the bottom.

**Extended Data Fig. 4.**
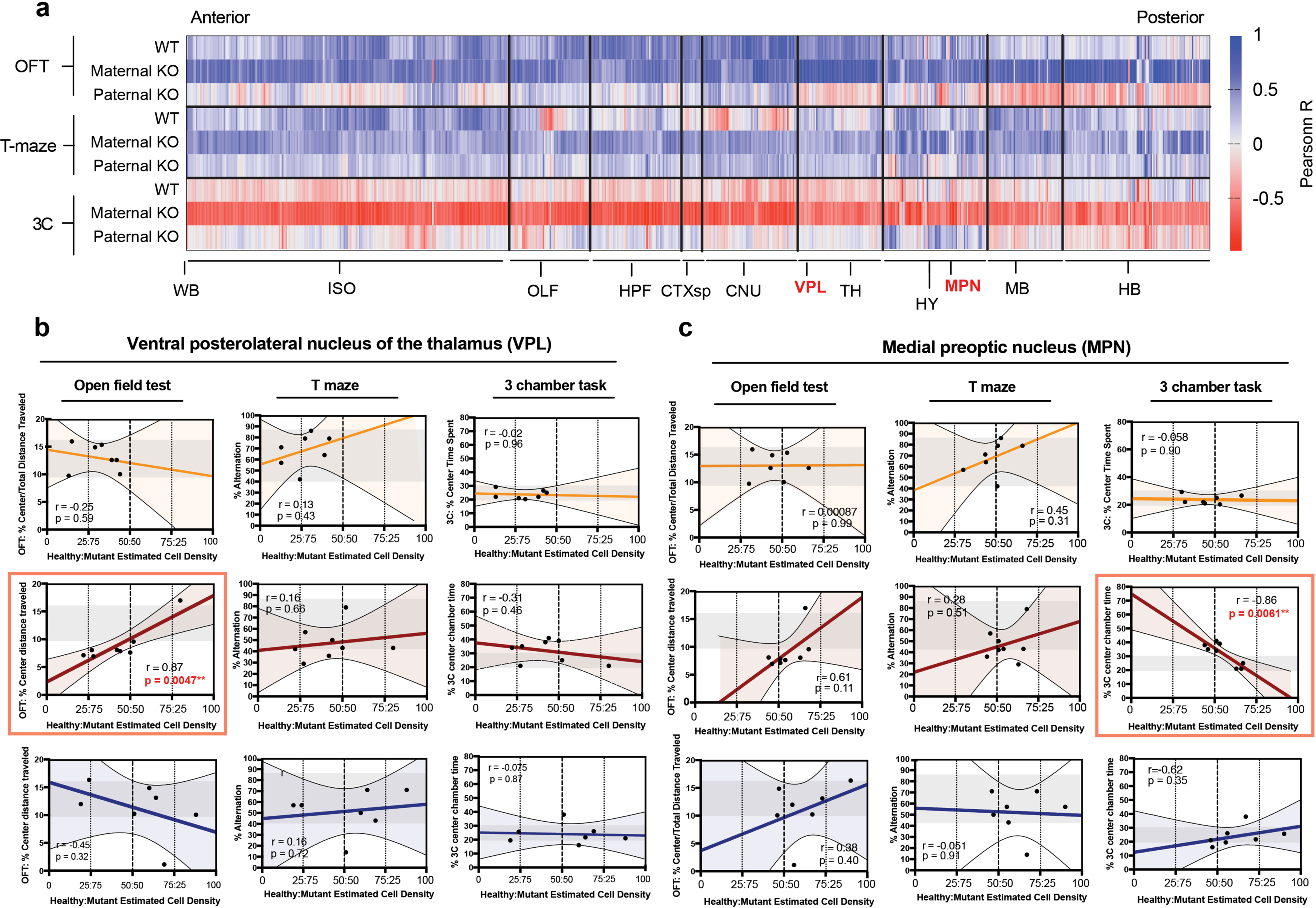
Identification of two sets of brain regions with local Fmr1-WT:KO cell ratios linked to phenotypes in the OFT and 3-chamber test: Raw Pearson r values and scatterplot examples across genetic groups and behavioral tests. **a**, 2D heat maps of raw Pearson correlation r values corresponding to **Fig 4a** across brain ROIs grouped by major ontological structures from anterior (left) to posterior (right) positions of the whole-brain. Raw Pearson r values are represented on a color gradient scale from -1 (red) to 0 (white) to 1 (blue). Results from Fmr1 WT (“WT”), maternal Fmr1-KO(m)/Mecp2-GFP(p) (“Maternal KO”) and paternal Fmr1-KO(p)/Mecp2-GFP(m) (“Paternal KO”) mice are grouped by each behavioral test: top: OFT, middle: T maze, bottom: 3-chamber (3C). **b**-**c**, Individual scatterplot visualization of the data from (**a**) for each behavioral group and behavioral test for the (**b**) ventral posterolateral nucleus of the thalamus (VPL), and (**c**) medial preoptic nucleus (MPN). Raw r and p values of statistical tests are inlayed within each panel. Red boxes bound correlations that reached statistical significance.

**Extended Data Fig. 5.**
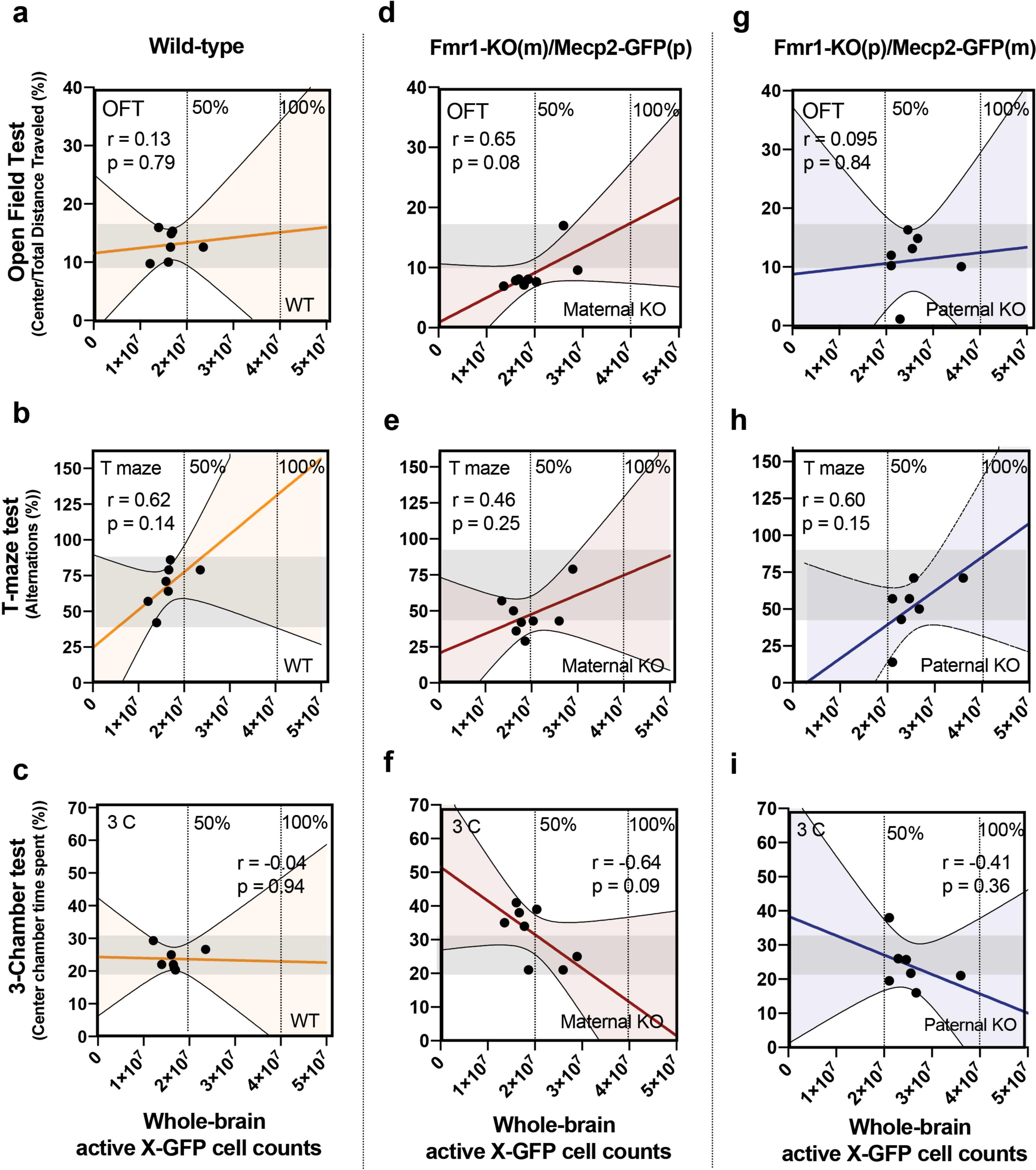
Whole-brain cell count correlations with behavior. (**a**-**c**; left) Fmr1 WT, (**d**-**f**; center) maternal Fmr1 KO, and (**g**-**i**; right) paternal Fmr1 KO whole-brain cell counts correlated to (**a**, **d**, **g**; top) OFT, (**b**, **e**, **h**; middle) T maze, and (**c**, **f**, **i**; bottom) 3 chamber behavioral scores. Dashed lines indicate estimated 50 and 100% cell counts. Gray transparent boxes inside plots indicate Fmr1 WT range of behavioral scores for comparison across mutant groups. r and p value statistics are listed in each panel.

**Extended Data Fig. 6.**
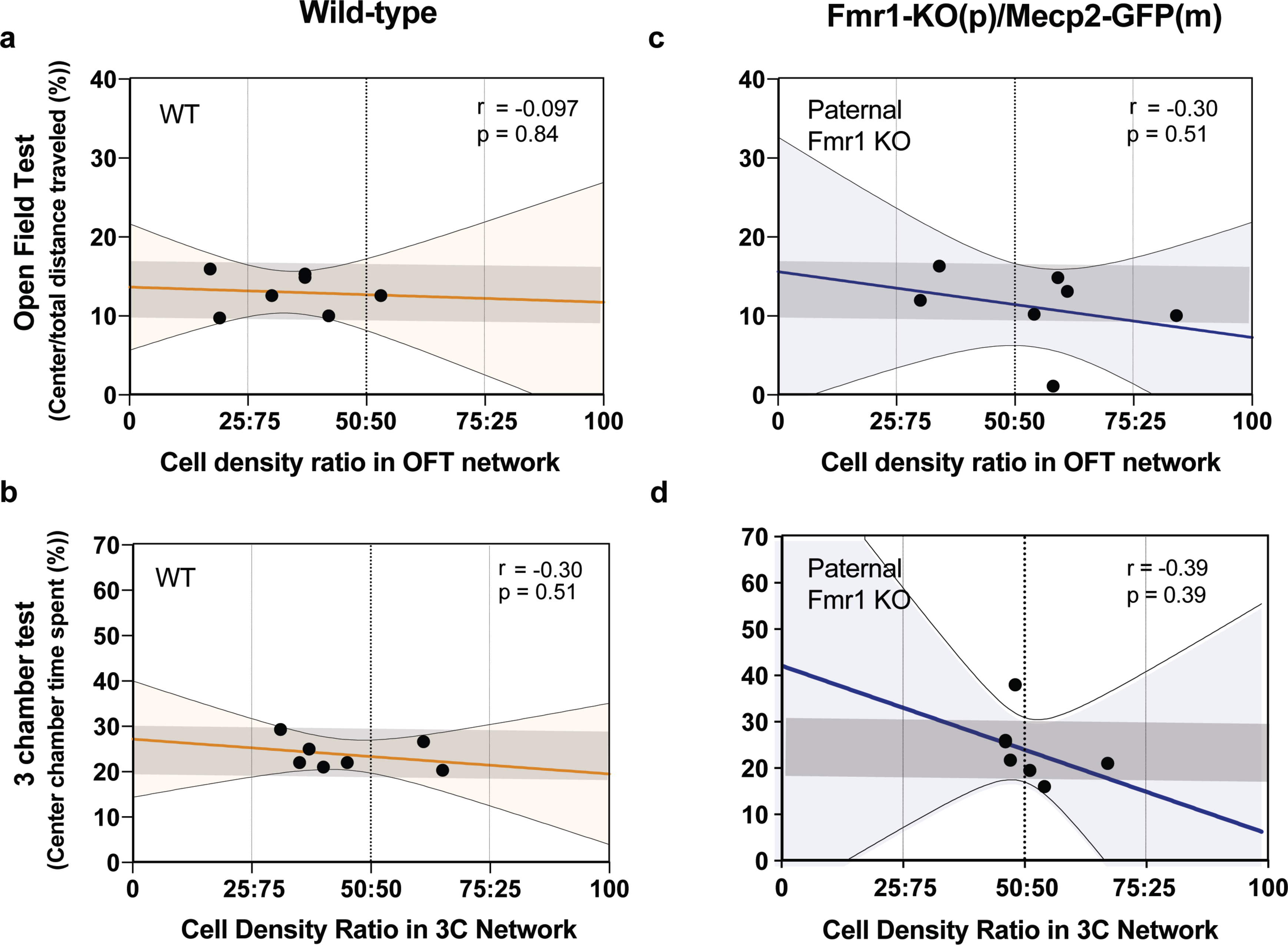
Control group correlations of behavioral brain network cell density ratios with behavioral scores. Fmr1 WT (**a**, **b**; left) and paternal Fmr1 KO (**c**, **d**; right) linear regression models of ROI network cell density ratios and scores from (**a**, **c**; top) OFT and (**b**, **d**; bottom) 3C behavioral assays. Regression and statistical test values are listed for each panel.

## Notes

https://github.com/AllenInstitute/chromosome-network-modeling

### Competing Interest Statement

The authors have declared no competing interest.

### Summary of Updates

This version of the manuscript includes a thorough revision of main text and figures throughout.

